# Dynamics of glioma-associated microglia and macrophages reveals their divergent roles in the immune response of brain

**DOI:** 10.1101/2021.07.11.451874

**Authors:** Jiawen Qian, Chen Wang, Bo Wang, Congwen Li, Kaiyi Fu, Yuedi Wang, Yiyuan Gao, Congyi Niu, Chujun Zhao, Jun Liu, Luman Wang, Ronghua Liu, Feifei Luo, Mingfang Lu, Yun Liu, Yiwei Chu

**Author notes:** These authors contributed equally to this work (Jiawen Qian, Chen Wang).

## Abstract

Glioma microenvironment contains numerous myeloid cells, including brain-resident microglia and recruited monocytes and macrophages (Mo/Mφ). When studied collectively, these cells presented pro-tumor effects. Yet, little is known about the differences among these myeloid populations. Using single-cell sequencing analysis, we studied the phenotypic characteristics, spatial variances, and dynamic changes of these relatively heterogeneous cell populations. Microglia populations with distinct spatial distribution presented different functional states, including tumor-associated subsets with phagocytic and lipid metabolism signature. Notably, this subset of glioma-associated microglia shared similar trait in a diverse spectrum of neuropathogenesis. In contrast, Mo/Mφ highly expressed genes related to angiogenesis, tumor invasion, and immune evasion. Moreover, identifying the Mo/Mφ subsets had prognostic and classificatory value in clinical application. These results thus eliminate the long-existing ambiguity about the role of microglia and Mo/Mφ in glioma pathogenesis, and reveal their prognostic and therapeutic value for glioma patients.

## Introduction

Glioma, a fatal malignancy in the CNS, develops highly complex immune microenvironment along with progression. Microglia and macrophages are the dominant immune cell subsets, composing approximately 30 - 50% of the cellular component in the tumor tissue(*1*). Most prior studies collectively termed these cells as glioma-associated microglia/macrophages (GAM) and revealed their multiple tumor-promoting effects, such as supporting tumor growth, angiogenesis, invasion, and immune evasion(*2, 3*). Therefore, GAM served as a theoretical target for glioma therapy. Unfortunately, blocking GAM activation with minocycline showed no clear clinical benefit, neither did GAM depletion with CSF-1R inhibitor(*2*). To improve the GAM-targeted therapy for glioma, more detailed studies are urgently needed to define the functional organization of GAM populations in the scenario of glioma.

GAM composed of two cell populations with distinct ontogeny. Microglia are the brain-resident phagocytes that derived from yolk sac, colonizing the CNS during the embryonic development and self-renewing instead of relying on peripheral supplements(*4*). Monocytes and macrophages (Mo/Mφ) entering afterwards originate from hemopoietic stem cell in the bone marrow, and gradually differentiate into mononuclear phagocytes during infiltration(*5*). The difference between microglia and Mo/Mφ have been explored in previous studies using microarray and bulk seq analysis(*6-8*). Recently, several researches unveiled the additional heterogeneity within these two cell populations based on the single-cell RNA sequencing (scRNA-seq) and mass cytometry (CyTOF) analysis(*9-15*). Exploring GAM cellular components may elucidate their potential functional differences in immune response. However, the depiction of microglia and Mo/Mφ subpopulations is still up for discussion. Their cellular component at different disease stage, their functional transition during glioma progression, and their spatial position in the tumor structure remained unknown.

Combining scRNA-seq analysis, microglia fate-mapping system, and multi-dimensional results, we depicted the temporal and spatial changes of microglia and Mo/Mφ during the development of glioma, and illustrated their phenotypic difference. According to previous research(*9, 10*) and our results, the majority of microglia located at the periphery of tumor foci. To avoid losing these microglial subsets, we thus isolated the myeloid cells from the tumor-bearing hemispheres, but not only the tumor mass. Based on this study, Mo/Mφ are more likely the factual mediator of tumor-promoting processes such as angiogenesis and T cells apoptosis. Microglia were misconceived as the saboteur to anti-tumor immune response. Instead, they may well be the sustainer of homeostasis in the CNS and the protector of neural tissue from the hyperactivated immune response. The expression pattern of *Cst7*^+^ microglia was common in multiple CNS pathologies. Moreover, significant correlation was found between particular Mo/Mφ subsets and the survival of glioma patients, indicating the predictive significance of these cell subsets.

## Results

### Different infiltration and spatial distribution between microglia and peripheral-derived monocytes/macrophages in murine glioma

To explore the dynamics of glioma-infiltrating microglia and Mo/Mφ, we established the orthotopic murine glioma model (Fig. 1A). The microglial fate-mapping system or reporter mice (*Cx3cr1*^*Cre*ER^: *R26*-*tdTomato*)(*16, 17*) were used to distinguish the microglia and Mo/Mφ populations (Fig. 1B). After tamoxifen treatment, microglia in the reporter mice were specifically labeled with red fluorescence, and can thus be stably and consistently traced, which was confirmed by flow cytometry analysis (Fig. 1C). Similarly, immunofluorescence staining also showed that microglia in the brain parenchyma were tdTomato^+^, while the meningeal macrophages derived from monocytes were free of the reporter protein (fig. S1A). These results showed that the fate-mapping mice model can properly distinguish the brain-resident microglia from peripheral-derived macrophages.

**Fig. 1.**
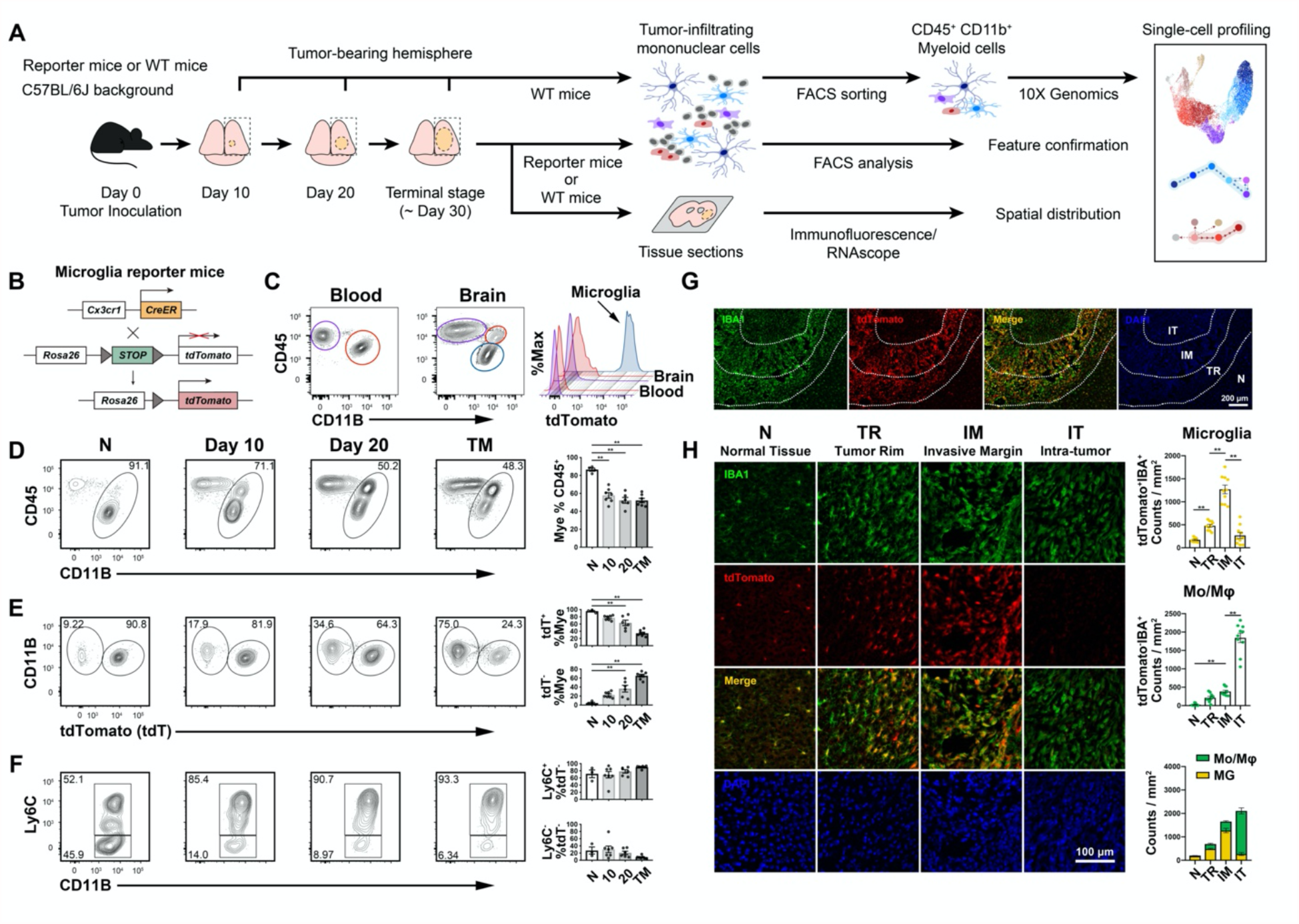
Infiltration and spatial distribution of myeloid cells in glioma. **(A)** Overview of the experimental strategy. **(B)** Schematic of *Cx3cr1*^CreER^:*Rosa26-tdTomato* microglia reporter mice. (C) FACS of the tdTomato expression in immune cell populations derived from glioma model (Day 20). (**D-F**) Left, FACS of myeloid cells in tumor-bearing hemispheres. Representative data of CD45^hi/lo^CD11^+^ Myeloid cells (**D**), tdTomato^+^ microglia and tdTomato^-^ Mo/Mφ (**E**), Ly6C^+^ Mo/Mφ and Ly6C^-^ Mo/Mφ (**F**) at indicated time points. Right, the statistical summary for abundance of the cell populations, n = 3-4 in normal tissue group, n = 6-7 in Day 10, Day 20, and Terminal Stage groups. Mye, CD45^hi/lo^CD11^+^ Myeloid cells. tdT, tdTomato. **(G)** Representative staining for IBA1 (green), tdTomato (red) and DAPI (blue) in the tumor-bearing hemispheres (Day 20). N, normal tissue; TR, tumor rim; IM, invasive margin; IT, intratumoral region. Dash line, tumor fraction border. Scale bar, 200 μm. **(H)** Left, Representative IF staining at indicated area of murine glioma (Day 20). Scale bar, 100 μm. Right, tdTomato^+^IBA1^+^ microglia or tdTomato^-^IBA1^+^ Mo/Mφ counts per field were calculated. Data were collected from 3-4 random fields for each region per mouse, n = 3. One-way ANOVA was performed in (**D-F**) and (**H**). * *P* < 0.05; * * *P* < 0.01. All values are shown as mean ± SEM.

Using flow cytometry, we first studied the proportion of myeloid cells infiltrated in the tumor-bearing brain. The SSC^hi^ granulocytes were excluded in all FACS analyses (fig. S1B). CD45^hi^ peripheral immune cells infiltrated CNS during tumor progression (fig. S1C). CD45^hi/lo^CD11b^+^ myeloid cells served as the majority of infiltrating immune cells and the cell number increased along with the tumor progression (Fig. 1D and fig. S1D). The proportion of tdTomato^-^ Mo/Mφ increased dramatically after Day 20 (Fig. 1E), in concordance with rapid tumor development(*18*). tdTomato^-^ subset primarily consisted of Ly6C^+^ cells (Fig. 1F). These results showed the dynamics of myeloid cell infiltration during the tumor progression.

Using Immunofluorescence staining, we found that while the amebiform IBA1^+^ tdTomato^-^ peripheral-derived Mo/Mφ accumulated in the core region of tumor foci (intratumoral region, IT), microglia mainly gathered at the periphery (Fig. 1G, H). As the microglia infiltrated from normal tissue (N) to invasive margin (IM), their morphology changed from typical ramified form to amoeboid and rod-like shapes (Fig. 1H). These results demonstrated that microglia and Mo/Mφ possessed distinct spatial distribution in the glioma microenvironment.

### scRNA-profiling of myeloid cells in glioma

To further investigate the phenotypic difference of the myeloid populations, we performed scRNA-seq for the CD45^hi/lo^CD11b^+^ cells isolated from the tumor-bearing hemispheres of the mice model at different stages of tumor development (Day 10, Day 20, Terminal Stage) (Fig. 1A). After filtering the low-quality cells and cells not in the study scope (NK cells, T cells, mast cells, and neutrophils), a total of 15,765 single-cell RNA profiles was used in following analyses. Graph-based clustering of the filtered myeloid cells identified 12 cell clusters (Fig. 2A), all of which expressed the pan-myeloid markers(*6-8*) (Fig. 2B). MG0-MG5 were featured with high expression of canonical microglia genes(*19, 20*) (Fig. 2C). In contrast, Mo1-2 and MP1-2 highly expressed canonical Mo/Mφ markers(*6-8*) (Fig. 2D). To verify the identity of various clusters, we assessed the microglia and Mo/Mφ identity score using gene set variation analysis (GSVA)(*21*) based on the reference genes of microglia (*P2ry12, Siglech, Sparc, Gpr34, Tmem119, Fcrls*, and *Olfml3*)(*8, 19, 20*) and Mo/Mφ (*Tgfbi, Ccr2, Ly6c2, Plac8, Itga4, S100a4*, and *S100a6*)(*6-8*) (Fig. 2E). Cell clusters were classified as either microglia or Mo/Mφ based on the identity score, and the results were confirmed using the expression of canonical markers respectively. As the main focus of the present study lies on microglia and Mo/Mφ, therefore, CD11b^-^ dendritic cells (mostly lost in the cell enrichment procedure) and CD11b^+^ dendritic cells were not included in further analysis. Similar to the flow cytometry results (Fig. 1E), the proportion of Mo/Mφ in the myeloid cells gradually increased during the tumor development (Fig. 2F, G). Some clusters, such as MG3, MG4, and MP2 also increased as tumor progressed, indicating their close relationship with tumor progression (Fig. 2H, I). In summary, the microglia and Mo/Mφ populations were distinguished according to the analysis of transcriptomic profile.

**Fig. 2.**
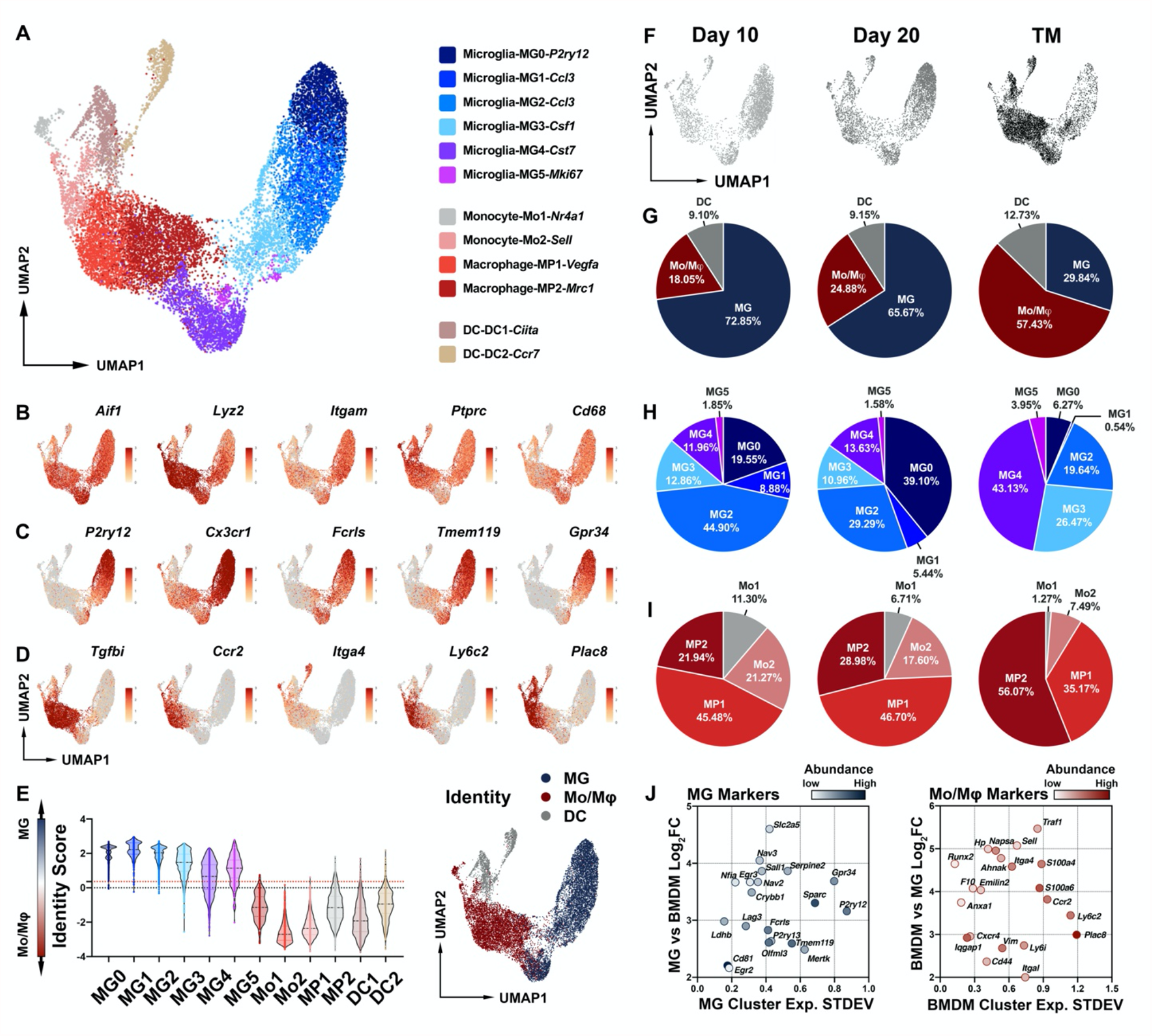
scRNA-seq profiling of myeloid cells in glioma. (**A**) UMAP projection showing 15,765 single-cell RNA profiles in 12 myeloid clusters. FACS-sorted CD45^lo/hi^CD11b^+^ myeloid cells were isolated from glioma model at Day 10, Day 20, and Terminal stage. Each sample of indicated time point was pooled from 5 mice (tumor-bearing hemispheres). (**B-D**) UMAP projection showing the expression of pan-myeloid (**B**), microglia (**C**), and Mo/Mφ **(D)**markers. **(E)** Left, violin plot representing the identity score of each cluster. Higher identity score indicated more “microglia-like”, and lower indicated more “Mo/Mφ-like”. Right, UMAP projection presenting the cell identity as microglia (MG), Mo/Mφ or dendritic cells (DC). **(F)** UMAP projection showing the myeloid cells at the indicated time points. **(G)** Pie plots representing the abundance of microglia (MG), Mo/Mφ, and dendritic cells (DC) at the indicated time points. (**H-I**) Pie plots showing the proportion of microglia (**H**) and Mo/Mφ (**I**) clusters at the indicated time points. (**J)** Scatter plots representing the robustness of microglia and Mo/Mφ markers in the glioma model. X axis, the standard deviation of mean expression of genes in the microglia (left) or Mo/Mφ (right) clusters. Y axis, the Log_2_ (Fold change) of mean expression of genes in the indicated populations. Shade of color, abundance of genes in the indicated populations.

Based on the general identity of each subgroup, we then investigated the expression of marker genes in different microglia and Mo/Mφ clusters. Notably, expression of some marker genes changed significantly during cell differentiation or activation, such as *Ly6C2* in Mo/Mφ and *P2ry12* in microglia (Fig. 2C, D). Furthermore, upregulation of typical microglia markers was found in some Mo/Mφ clusters, including *Olfml3, Fcrls* and *Tmem119* (fig. S2A). Therefore, we verified the stability and robustness of marker genes within the two subgroups, *Crybb1* and *Ldhb* were expressed abundantly, specifically, and consistently in microglia clusters, while *Iqgap1* was recognized as a robust cell marker for Mo/Mφ clusters (Fig. 2J and fig. S2B). According to the Immunological Genome Project (ImmGen ULI RNA-seq) dataset(*22*), microglia rarely express *Iqgap1* compared with many myeloid cell subsets (fig. S2C). Confirmed in Ivy Glioblastoma Atlas Project (IVYGAP) dataset(*23*) (fig. S2D) and the GBM single-cell sequencing dataset (GSE84465)(*9*) (fig. S4E, H), *IQGAP1* can also distinguish Mo/Mφ from microglia in human glioma samples. Tissue immunohistochemistry results from the Human Protein Atlas database(*24*) indicated that IQGAP1 mainly located at the perivascular space in normal brain, and its distribution pattern was completely different from microglia marker P2RY12 (fig. S2E). We have thus identified stable gene markers to distinguish microglia and Mo/Mφ, which is of great significance to the study of myeloid cells in CNS.

### Phenotypical transition, temporal changes and spatial distribution of microglia during the progression of glioma

We then preformed differentially expressed gene (DEG) analysis and examined the transcriptional status of microglial clusters (Fig. 3A, C, D and Table S2). Cell trajectory analysis depicted the linear continuum of microglia clusters (Fig. 3B). MG0 was recognized as the “resting” microglia in the normal tissue, since this cluster was featured with expression of genes related to the homeostasis of CNS, including *P2ry12, Gpr34*, and *Smad7*(*25-27*) (Fig. 3C-F). MG1-2 were pro-inflammatory microglia, considering the expression of inflammatory factors (*Tnf*) and chemokines (*Ccl3* and *Ccl4*) (Fig. 3C, D, F). Interestingly, MG3 and MG4 shared similar transcriptomic features with Mo/Mφ. As previously reported, glioma-associated myeloid cell markers *Spp1* were found highly expressed in these two clusters(*6*), suggesting their correlation with glioma scenario (Fig. 3C and Fig. 4F). Furthermore, phagocytosis-associated gene *Cst7* was also a feature gene of MG3 and MG4(*28*) (Fig. 3C, D, F). Moreover, genes involved in fatty acid metabolism such as *Fabp5* presented high expression in these two clusters(*29*), suggesting their unique metabolic feature (Fig. 3C, D). MG5, expressing proliferation-associated genes (*Mki67, Cenpf, Cdk1, Birc5, Stmn1* and *Top2a*), were actively proliferating microglia (Fig. 3C, D and fig. S3A), suggesting that microglia accumulated at tumor periphery derived from both migration and proliferation.

**Fig. 3.**
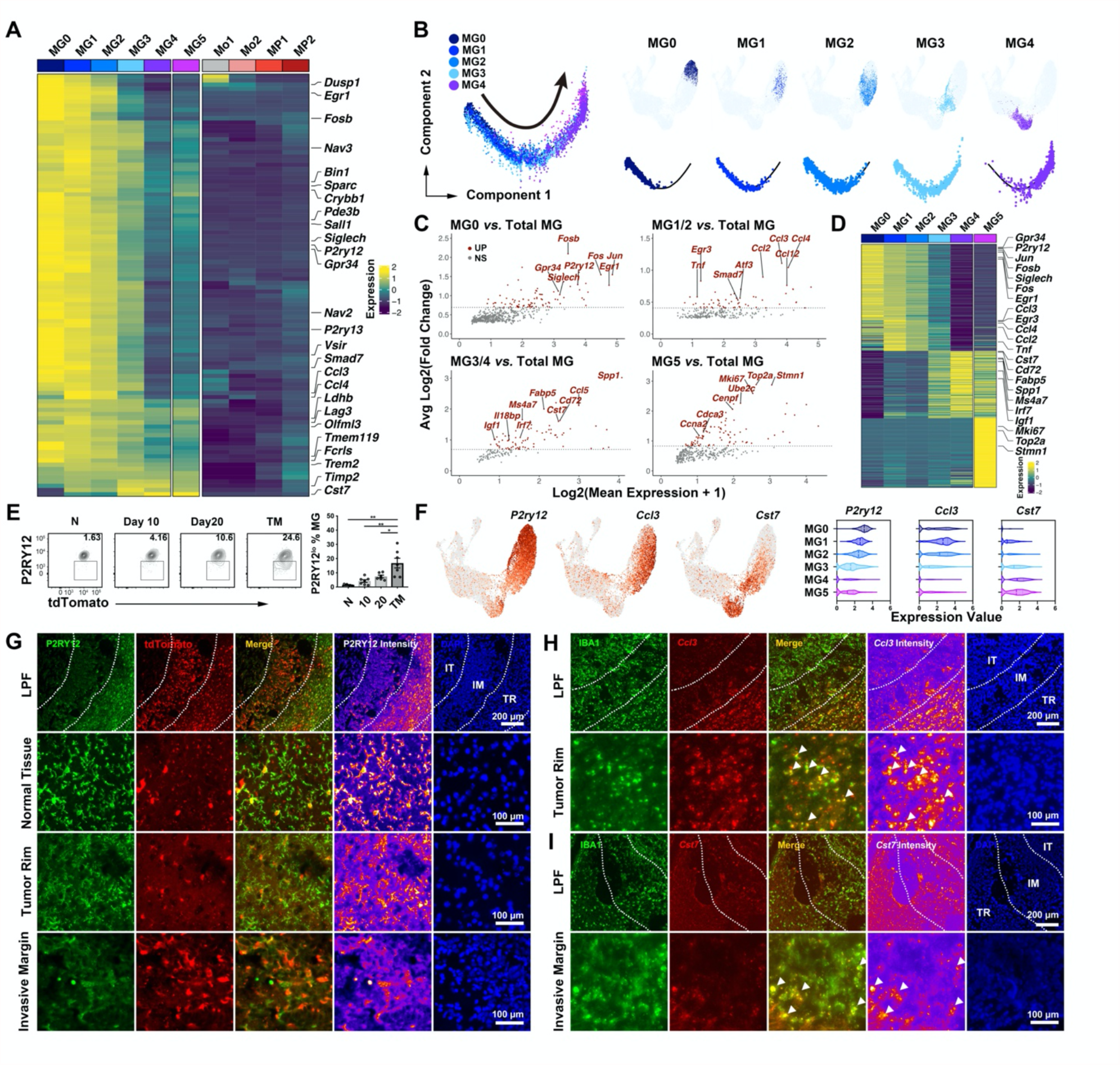
Microglial components in glioma. **(A)** Heatmap indicating the expression of microglia enriched genes. **(B)** Left, pseudotime-ordered analysis of microglial clusters in glioma. Right, UMAP projection and 2D pseudotime plots of each microglial subset. **(C)** MA plots displaying genes that are upregulated (red) in each microglial cluster. Dashed lines denote fold change thresholds used when identifying DEGs. **(D)** Heatmap indicating the expression of marker genes for microglial clusters. **(E)** Left, flow cytometry analysis of P2RY12 expression in tdTomato^+^ microglia (MG). Right, the statistical summary for abundance of P2RY12^+^ microglia, n = 3-4 in normal tissue (N) group, n = 6-7 in Day 10 (10), Day 20 (20), and Terminal Stage (TM) group. One-way ANOVA was performed. * *P* < 0.05; * * *P* < 0.01. All values are shown as mean ± SEM. **(F)** Left, UMAP projection showing the expression of selected marker genes for microglial clusters. Right, violin plots representing the expression of marker genes in each cluster. (**G-I**) Representative images of IF staining for P2RY12 in glioma at Terminal Stage (**G**), RNAscope for *Ccl3* at Day 10 (**H**) and *Cst7* at Day 20 (**I**). Fields from low power field (LPF) and indicated area of murine glioma were shown. TR, tumor rim; IM, invasive margin; IT, intratumor. Dashed line, border of different areas. Arrows, *Ccl3*^*+*^ (**H**) or *Cst7*^*+*^ (**I**) microglia. Scale bar, 200/100 μm.

**Fig. 4.**
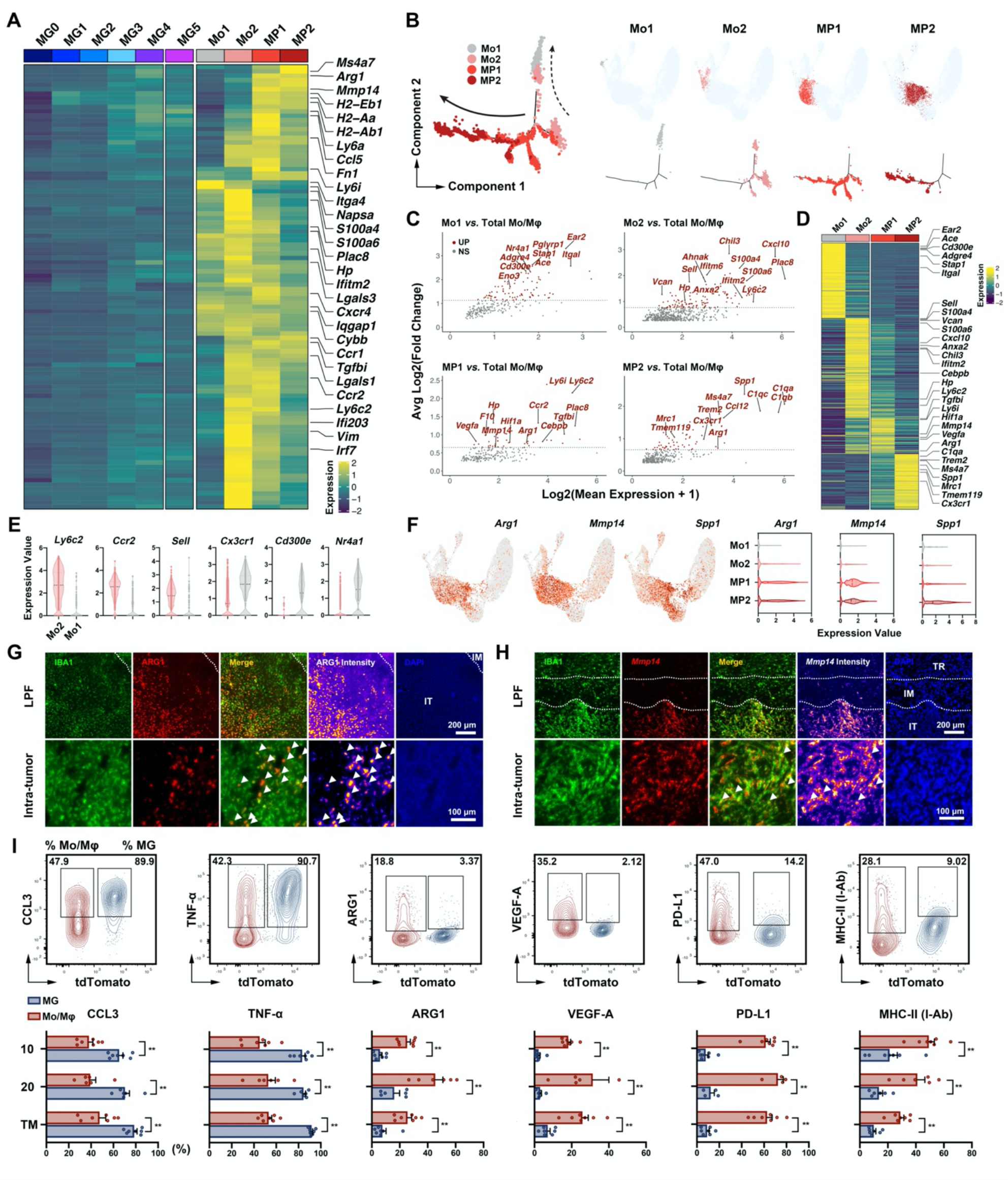
Monocyte/macrophage components in glioma. **(A)** Heatmap indicating the expression of Mo/Mφ enriched genes. **(B)** Left, pseudotime-ordered analysis of Mo/Mφ clusters in glioma. Right, UMAP projection and 2D pseudotime plots of each microglial subset. **(C)** MA plots displaying genes that are upregulated (red) in each Mo/Mφ cluster. Dashed lines denote fold change thresholds used when identifying DEGs. **(D)** Heatmap indicating the expression of marker genes for Mo/Mφ clusters. **(E)** Violin plots showing the expression of featured genes involved in the transition from Mo2 (Ly6C^+^) to Mo1 (Ly6C^-^). **(F)** Left, UMAP projection showing the expression of selected marker genes for Mo/Mφ clusters. Right, violin plots representing the expression of marker genes in each cluster. (**G-H**) Representative images of IF staining for ARG1 at Terminal Stage (**G**) and RNAscope for *Mmp14* at Day 20 (**H**). Fields from low power field (LPF) and indicated area of murine glioma were shown. TR, tumor rim; IM, invasive margin; IT, intratumoral region. Dashed line, border of different areas. Arrows, ARG1^+^ (**G**) or *Mmp14*^+^ (**H**) myeloid cells. Scale bar, 200/100 μm. (**I**) Top, flow cytometry analysis of featured markers in microglia (MG, tdTomato^+^) and Mo/Mφ (tdTomato ^-^). Bottom, the statistical summary for expression of featured markers, n = 4-7. Student’s t-test was performed, * *P* < 0.05; * * *P* < 0.01. All values are shown as mean ± SEM.

The distribution of feature genes was consistent with the microglial trajectory (fig. S3C). The transition of microglia started from “resting” state to activated and pro-inflammatory state, and finally reached an end stage of tumor-associated phenotype. We also checked the abundance of microglia clusters at different stages of glioma model (Fig. 2H and Table S1). As expected, the majority of the microglia presented pro-inflammatory state (MG1-2) at the early stage (Day 10) of glioma development, and the majority of the microglia presented glioma-associated phenotype (MG3-4) at the terminal stage (TM) (Fig. 2H).

Immunofluorescence and RNAscope were performed to validate the distribution of microglia subsets in the glioma microenvironment. As expected, tdTomato^+^ microglia in the normal tissue possessed higher expression of P2RY12 compared with those at the tumor rim, and hardly any expression of P2RY12 was observed in the microglia at the invasive margin (Fig. 3G). The expression pattern of P2RY12 in microglia validated the previous analysis of cluster identity and phenotype. Flow cytometry analysis showed that P2RY12^lo^ tumor-associated microglia gradually increased as tumor progressed (Fig. 3E). RNAscope results indicated that *Ccl3*^+^ pro-inflammatory MG1-2 located at the tumor rim (Fig. 3H), *Cst7*^+^ glioma-associated phenotype mainly gathered at the invasive margin and intra-tumoral area (Fig. 3I), and *Mki67*^+^ proliferating microglia were found at invasive margin (fig. S3B). Collectively, these analyses revealed the phenotype transition, temporal changes and spatial distribution of microglia during the progression of glioma.

### Phenotypic transition, temporal changes and spatial distribution of Mo/Mφ during the progression of glioma

We then analyzed the tumor-infiltrating Mo/Mφ, unsupervised clustering identified 4 Mo/Mφ clusters in our dataset and feature genes for each cluster were identified by DEG analysis (Fig. 4A, C, D and Table S2). Cell trajectory analysis depicted a branched structure of Mo/Mφ clusters (Fig. 4B). Mo1 was featured with high expression of peripheral monocyte markers (*Ace, Ear2*, and *Cd300e*) and low expression of *Ly6c2*(*30-32*) (Fig. 4C, D). Mo2, on the contrary, highly expressed *Ly6c2* and various mononuclear phagocyte markers (*Chil3, Hp, Vcan*, and *Sell*)(*30-32*) (Fig. 4C, D, E), suggesting that this cluster was the Ly6C^+^ monocytes that skewed to differentiate into macrophages. In concordance with the transition of peripheral monocytes reported in recent studies(*30-32*), Mo1 were recognized as Ly6c^-^ monocytes, since this cluster manifested comparatively low level of *Ly6c2, Ccr2* and *Sell* as well as high expression of *Cx3cr1, Cd300e* and *Nr4a1* (Fig. 4E). Considering the specific expression of *Arg1* and *Mmp14* in MP1 and MP2 (Fig. 4C, D, F), these two clusters were most probably the accomplice to tumor progression. These two molecules played a critical role in promoting tumor progression and were reported to have common expression in the tumor-infiltrating myeloid cells(*2, 3*). ARG1 was found to mediate the inhibition of tumor-infiltrating T cells(*33*), while MMP14 was responsible for the invasive growth of glioma(*34, 35*). These two genes were expressed exclusively in Mo/Mφ, suggesting that microglia were the scapegoat of tumor-promoting phenotype of Mo/Mφ. Moreover, MP1 highly expressed *Vegfa*, closely related to the tumor angiogenesis(*36*) (Fig. 4C, D). Besides, MP2 differentially expressed *Mrc1* (Fig. 4C, D), suggesting that these cells were related to angiogenesis, tumor invasion, and immune suppression(*37*). Immunofluorescence staining and RNAscope results verified the main distribution of ARG1^+^ and *Mmp14*^+^ MP1-2 in the intra-tumoral region (Fig. 4G, H).

Based on the known transition from Ly6C^hi^ monocytes to Ly6C^lo^ monocytes(*31*), Mo2 (Ly6C^hi^ monocytes) was set as the start of inferred trajectory. Mo2-MP1-MP2 transition appeared as a branched structure along the main trajectory (Fig. 4B). This was consistent with the distribution of feature genes along the trajectory (fig. S3D). This analysis demonstrated the differentiation of Mo/Mφ along with infiltration, as well as their eventual transition to the tumor-associated macrophage. Further, the abundance of Mo/Mφ clusters at the early and middle stage of glioma progression was similar, whereas massive increase of MP2 was observed at the terminal stage (Fig. 2G, I). Combined with the fulminant tumor progression observed at Day 20, MP2 likely played an indispensable role in the explosive growth of tumor. Flow cytometry analysis of featured markers in tdTomato^+^ microglia and tdTomato ^-^ Mo/Mφ further confirmed the differences between these populations (Fig. 4I). Conclusively, these analyses revealed that Mo/Mφ phenotype transition, temporal changes and spatial distribution of Mo/Mφ during the progression of glioma.

### Single-cell profiling of myeloid cells in human glioma sample

We then studied myeloid cells in human glioma sample. We re-clustered myeloid cells in human glioma dataset (data from GSE84465)(*9*), and identified 6 cell clusters (fig. S4A). C0 and C5 showed high expression of canonical microglia genes (*P2RY12* and *GPR34*) (fig. S4D), while C2 and C3 were featured with expression of Mo/Mφ markers (*TGFBI, S100A6* and *IQGAP1*) (fig. S4E). The corresponding identity score further verified the identity of cell clusters (fig. S4C). Microglia clusters in human shared high expression of microglia marker genes found in mice (*CCL3, CCL4*, and *TNF*) (fig. S4F). Consistent with their identities, C2 and C3 highly expressed Mo/Mφ marker genes (*TGFBI, VEGFA*, and *GPNMB*) (fig. S4E, G). Similar distribution pattern of microglia and Mo/Mφ clusters were found in human samples and mice model. Microglia gathered at the peripheral region while Mo/Mφ accumulated in the core area of tumor (fig. S4B). Similar distribution of microglia and Mo/Mφ was also confirmed in the IVYGAP dataset (fig. S4H). Interestingly, C1 and C4 were of mixed identity and presented both characteristics of microglia and Mo/Mφ (fig. S4C). This was possibly due to the prolonged course of glioma in human, during which Mo/Mφ adapted to the CNS microenvironment and acquired microglia-like phenotypes. In summary, myeloid cells in human glioma sample and mice glioma model presented similar transcriptome profile and distribution pattern.

### Pathway enrichment and functional transition of microglia and Mo/Mφ clusters

To investigate biological states and functional characteristics of various myeloid subsets, we analyzed the pathways enrichment in microglia and Mo/Mφ by GSVA. Gene sets were selected from Gene Ontology (GO) and Kyoto Encyclopedia of Genes and Genomes (KEGG) pathway database (Table S3). Pathways involved in activation and migration gradually increased during the transition of both microglia and Mo/Mφ, while enrichment of pathway related to antigen processing and presentation turned to decrease during the transition (Fig. 5A), in consistent with the previous study(*18*). Enrichment of pathway related to morphological changes (dendritic spine development) steadily decreased during the transition (Fig. 5A), consistent with the observation in immunofluorescence staining. Pathway analysis also revealed the different characteristics of microglia and Mφ subsets (Fig. 5B-H).

**Fig. 5.**
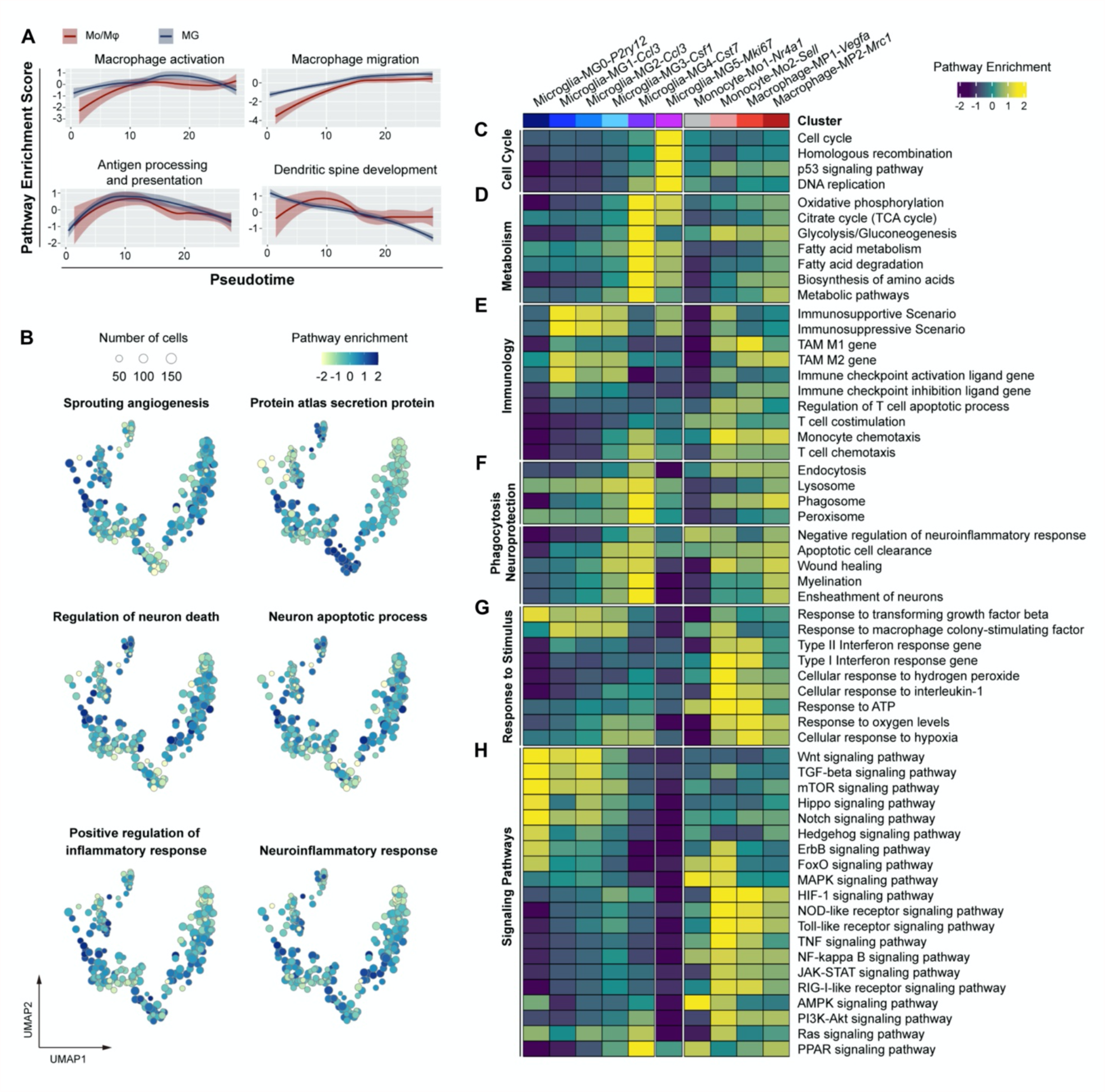
Pathway enrichment and functional transition of microglia and Mo/Mφ clusters. **(A)** Two-dimensional plots showing the pathway enrichment scores for genes related to macrophage activation, macrophage migration, antigen processing, and presentation and dendritic spine development in Mo/Mφ (red) and microglia (MG, blue) along with the pseudotime. **(B)** A representative UMAP projection showing the enrichment of pathways related to angiogenesis, inflammation response, neuroprotection and protein secretion. Size of the bubbles indicated the number of cells. (**C-H**) Heatmap indicating the enrichment of pathways related to cell cycle (**C**), metabolism (**D**), immunology (**E**), phagocytosis and neuroprotection (**F**), response to stimulus (**G**), and signaling pathways (**H**). GSVA was applied to calculate the pathway enrichment based on the mean expression of genes within each subset.

The enrichment of terms related to response to stimulus provided additional information about the spatial distribution (Fig. 5G). Mo1 were least enriched in transforming growth factor beta (TGF-β) related terms and hypoxia-related pathways (Fig. 5G, H), and most likely remained in the oxygen-rich blood vessels instead of entering the brain parenchyma that contains high concentration of TGF-β(*38*). As reported previously, Ly6C^-^ monocytes such as Mo1 cannot enter parenchymal tissues(*31, 32*). On the contrary, Mo2, MP1 and MP2 were comparatively enriched in terms related to inflammatory response and hypoxia-related pathways, suggesting that they existed in hypoxic intra-tumoral tissues (Fig. 5G, H). These results were consistent with the location distribution of myeloid populations observed by immunofluorescence.

Generally, activation process of microglia and Mo/Mφ were similar. Some pathway scores gradually increased during the transition of microglia and Mo/Mφ, including pathways involved in activation, migration, and phagocytosis (Fig. 5A), whereas other pathways only enriched transiently, including pathways involved in nerve damage (regulation of neuron death and neuron apoptotic process), pathways related to neuroinflammation (antigen processing and presentation and neuroinflammatory response), and various immune-related signaling pathways (TLR signaling, NF-κB signaling and JAK-STAT signaling) (Fig. 5A, B, H), consistent with our previous observations(*18*). Similar pattern was also found in human glioma-associated myeloid cells (fig. S5).

However, microglia and Mo/Mφ clusters eventually evolved to enrich in divergent functions. Enrichment score for M1/M2 phenotype changed synchronously in the transition of microglia clusters, while in Mo/Mφ clusters, enrichment score for M2 phenotype continued to increase even when M1 phenotype score decreased (Fig. 5E). These results suggested that pro-inflammatory effect of microglia was self-restricted, thereby preventing excessive inflammation in the CNS. As for Mo/Mφ, the M2-biased phenotype of MP2 may suppress the anti-tumor immune response. Additionally, MP1 and MP2 were highly enriched of terms involved in immune checkpoint and apoptosis process of T cells (Fig. 5E), suggesting that Mo/Mφ facilitated the immune evasion of glioma.

We then performed SCENIC analysis and obtained the featured regulons of each cluster (fig. S6A). AP-1 family members such as Jun and Fos were highly active in “resting” MG0 (fig. S6B). However, previous study proposed that cell isolation process of inactivated sensitive cells such as MG0 may lead to the up-regulation of these genes(*20*). Atf3 was highly active in MG1 and MG2, suggesting its pro-inflammatory role in these subsets (fig. S6B). Moreover, we found that Nfia might serve as a linage-related regulon to distinguish microglia from Mo/Mφ (fig. S6B). As for Mo/Mφ clusters, Cebpb was highly activated (fig. S6B). Irf1 and Irf7 were active in pro-inflammatory Mo2 and MP1 (fig. S6B).

### Disease-associated gene expression pattern of microglia

Among microglial clusters, MG4 was the most abundant cluster at the terminal stage of glioma and was the end of phenotype transition (Fig. 2H, 3B and Table S1). The transcriptome profile of MG4 was characteristic of active metabolism and protein secretion, strong phagocytic function and decreased immune activity (Fig. 5B, D-F). Surprisingly, such disease-associated pattern was also found in many CNS pathological processes. Analysis of terms related to various CNS diseases showed that corresponding enrichment score steadily increased during the transition of microglia and reached the peak value in MG4 (Fig. 6A, B). Then, we analyzed the identity of disease-associated myeloid cells based on marker genes of disease-associated subset (Table S4) and MG4 respectively. Either in Alzheimer disease model (5xFAD and CK-p25)(*39, 40*), aging mice(*41*) or demyelination model (experimental autoimmune encephalitis, EAE and lysolecithin injection, LPC)(*41, 42*), myeloid cells high in disease-associated microglia score coincided with those high in MG4 identity score (Fig. 6C). These results suggested that disease-associated transcriptome pattern found in MG4 might serve as the mutual characteristics of CNS pathologies.

**Fig. 6.**
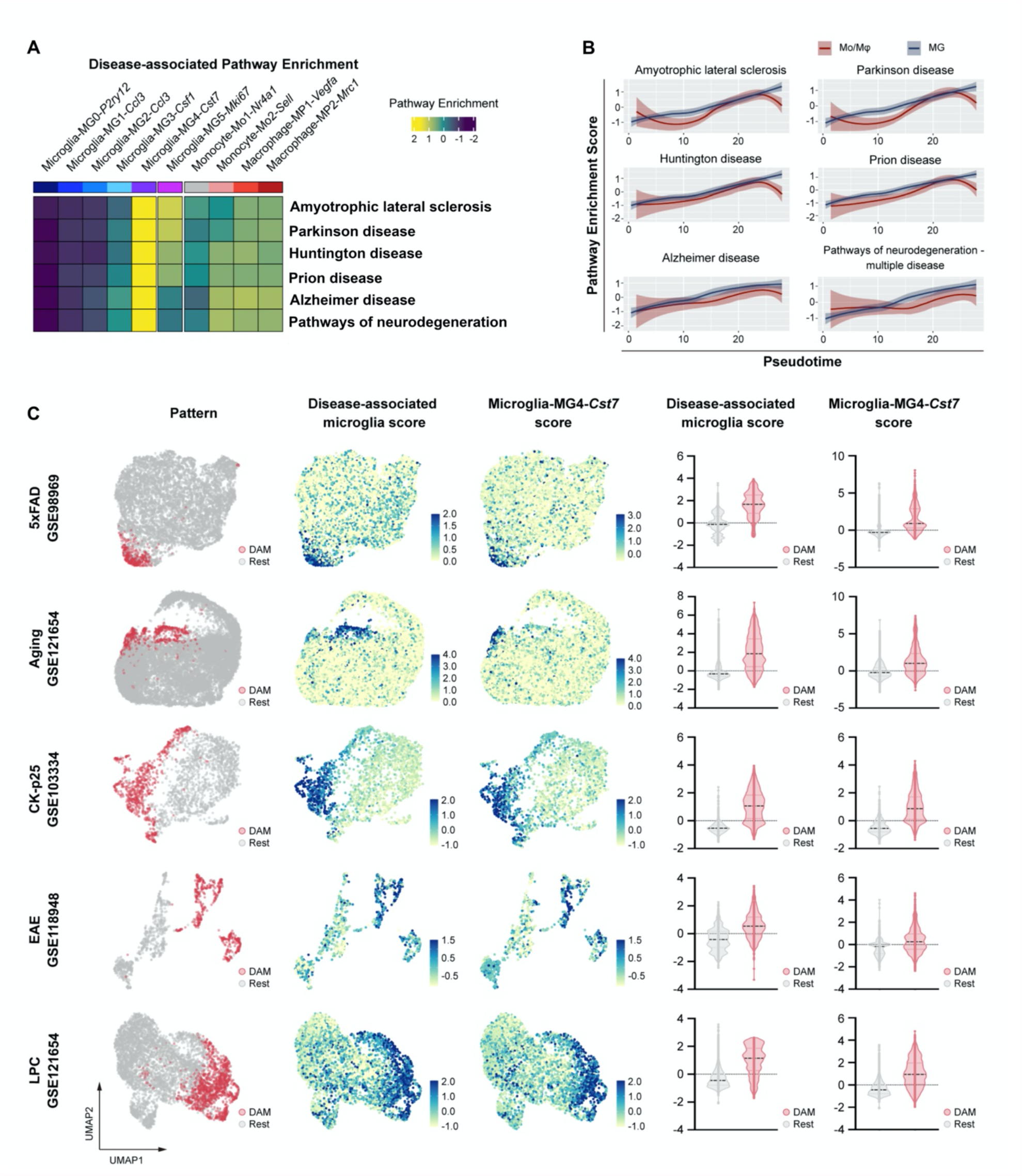
Common disease-associated pattern found in multiple CNS pathology. **(A)** Heatmap indicating the enrichment of disease-associated pathways. GSVA was applied to calculated the pathway enrichment based on the mean expression of genes within each subset. **(B)** Two-dimensional plots showing the enrichment scores for disease-associated pathways in Mo/Mφ (red) and microglia (MG, blue) along with the pseudotime. **(C)** Left, UMAP projections showing disease-associated microglia (DAM, red) and rest myeloid cells (Rest, grey), disease-associated microglia score, and microglia-MG4-*Cst7* score in myeloid cells from 5xFAD mice (data from GSE98969), aging mice (data from GSE121654), Ck-p25 mice (data from GSE103334), EAE mice (data from GSE118948), and LPC mice (data from GSE121654). Right, violin plots showing the disease-associated microglia score and microglia-MG4-*Cst7* score of disease-associated microglia (DAM, red) and rest myeloid cells (Rest, grey) for corresponding datasets. GSVA was applied to calculate the disease-associated microglia score and microglia-MG4-*Cst7* score. Marker gene lists of disease-associated microglia were concluded from the previous studies (Table S4).

### Composition of the TCGA LGG/GBM cohort and the clinical implication

We next explore the potential clinical applications of the myeloid cell composition using TCGA LGG/GBM dataset. Survival analysis showed that expression of microglia feature genes (*CCL3, P2RY12* and *PDE3B*) was positively correlated with prognosis of disease, but patients with high expression of Mo/Mφ feature genes (*IQGAP1, VEGFA, MMP14, ARG1* and *ITGA4*) presented unfavorable prognosis (fig. S7). We then estimated the cellular composition of glioma using MCP-counter. According to the infiltration of peripheral immune cells and dominant myeloid populations, TCGA LGG/GBM cohort was divided into 6 groups (Fig. 7A-C). Survival analysis indicated that patients with high infiltration of peripheral immune cells had prominently poor prognosis (Fig. 7D). Consistently, flow cytometry analysis revealed that peripherally infiltrated CD45^hi^ immune cells increased during tumor progression (fig. S1C). Furthermore, in groups with high infiltration of peripheral immune cells, median survival time (MST) of Mo/Mφ-dominant group (13.6 months) was much shorter than the MST of microglia-dominant group (47.9 months) (Fig. 7D). Same trend also existed in the median and low infiltration groups. Moreover, positive correlation was found between abundance of quiescent microglia and the prognosis of glioma, abundance of monocytes and macrophages were negatively correlated with patient prognosis (Fig. 7E).

**Fig. 7.**
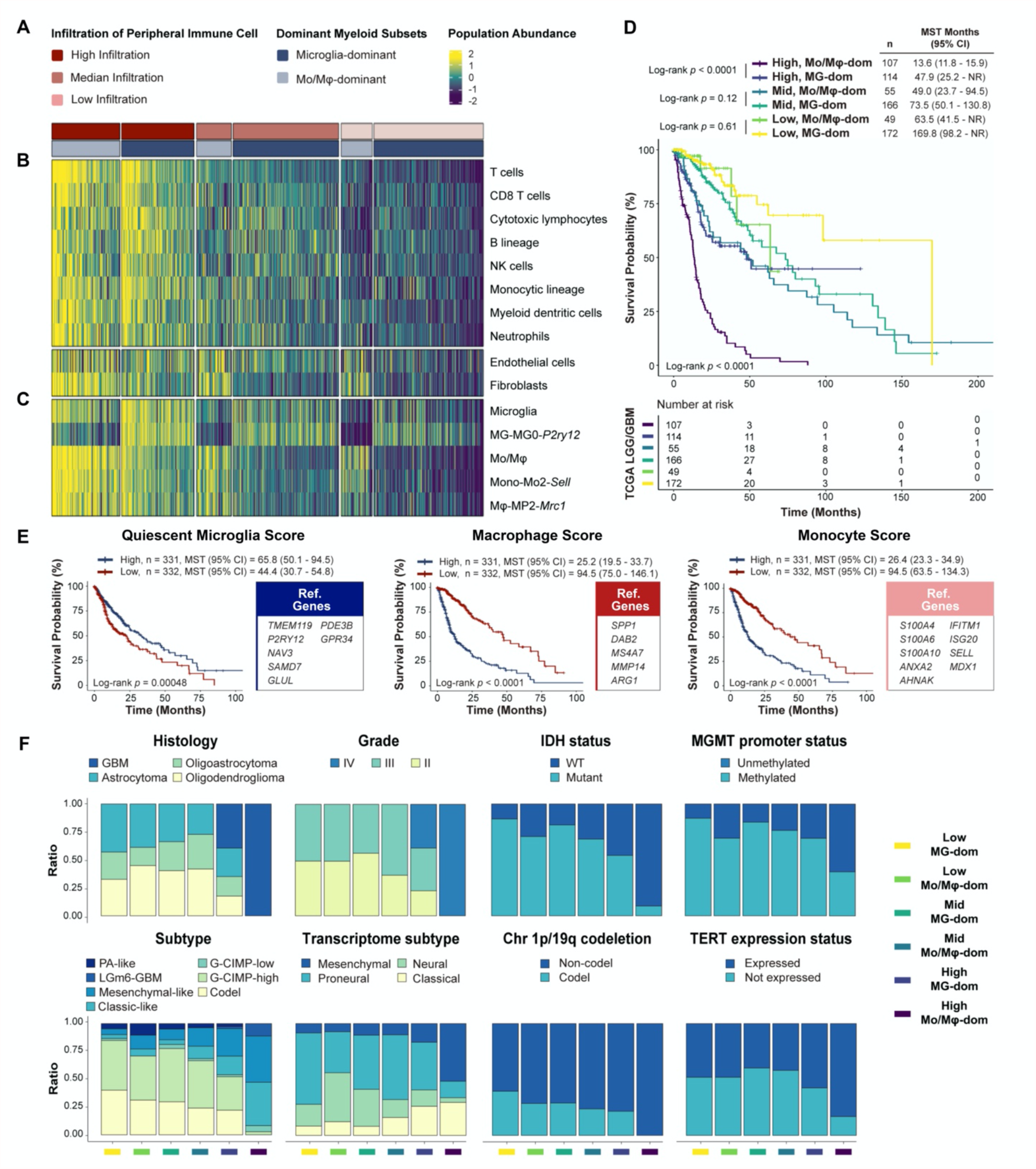
Composition of the TCGA LGG/GBM cohort and the clinical implication. **(A)** Composition of the TCGA LGG/GBM cohort by infiltration of peripheral immune cells and dominant myeloid cells. **(B)** Heatmap showing the composition of the tumor microenvironment defined by the MCP-counter Z-scores. **(C)** Heatmap showing the abundance scores for microglia, MG-MG0-*P2ry12*, Mo/Mφ, Mono-Mo2-*Sell* and Mφ-MP2-*Mrc1* subsets defined by the MCP-counter Z-scores. **(D)** Kaplan-Meier analysis showing the survival probability of patients in each composition of the TCGA LGG/GBM cohort. The number of patients and the risk classification are indicated in the figure. Significance was calculated using the log-rank test. High, Mid, or Low represented high, median, or low infiltration of peripheral immune cells. Mo/Mφ-dom, monocyte and macrophage dominant subgroup. MG-dom, microglia dominant subgroup. n, number of patients. MST, median survival time. **(E)** Left, Kaplan-Meier analysis showing the survival probability of patients, characterized by either high (blue) or low (red) abundance of quiescent microglia (left), macrophages (middle) and monocytes (right) subsets, respectively. Right, marker genes used to estimate the abundance of corresponding subset with MCP-counter. **(F)** Histograms showing the proportion of patients in various classifications of the TCGA LGG/GBM cohort.

Then, we examined the composition of patients in 6 groups according to various classification of LGG/GBM cohort(*43, 44*) (Fig. 7F). Isocitrate dehydrogenase (IDH) mutation, codeletion of the 1p and 19q chromosome arms (Chr 1p/19q), O6-methylguanine-DNA methyltransferase (MGMT) promoter methylation status and telomerase reverse transcriptase (TERT) expression and promoter status are all significant prognostic biomarkers for glioma patients. Subtype and transcriptome subtype are comprehensive classification based on the molecular and genetic profiles of glioma patients. We found that composition of patients in 6 groups was quite different based on the authoritative and comprehensive classification of glioma, and was consistent with the prognostic implications of these classifications (Fig. 7F). In conclusion, these results revealed the prognostic and classificatory value of Mo/Mφ. The newly proposed classification of glioma patients might serve as reference for prognosis prediction and target therapy.

## Discussion

It is of great interest to distinguish the glioma-associated myeloid populations, which have distinct origins and thus potential differences in phenotype and function. In this study, we leveraged a diverse panel of analyses and datasets to evaluate the brain-resident microglia and Mo/Mφ in the glioma microenvironment. Differences between these populations were found not only in spatio-temporal dynamics, but also in transcriptomic profiles, pathway enrichment, and prognosis implication. Microglia may well initiate anti-tumor immune response in the early phase of tumor development, but ultimately skew to phenotypes involved in phagocytic clearance, restoration of CNS homeostasis and tissue protection. Besides, a disease-associated microglia cluster was found universal in a various spectrum of neurological diseases. Mo/Mφ, on the other hand, gradually invaded and accumulated during tumor progression, most likely prohibited the anti-tumor immune response and promoted tumor angiogenesis and invasion. In conclusion, microglia have most probably been misconceived as the scapegoat of tumor-promoting Mo/Mφ.

The difference in spatial distribution between Mo/Mφ and microglia was first noticed in the bulk-seq for microdissection of specific glioma structure and scRNA-seq for tumor tissue of different region(*9, 10*). Antunes *et al*.(*15*) and we validated the distribution difference in animal models through the microglial fate-mapping system. In addition, Caponegro *et al*.(*45*) and we found that the activated *Ccl3*^+^ microglia accumulated in the tumor rim. This indicated that, compared with “clean tumor-bulk”, obtaining cell samples from the entire tumor-bearing hemisphere can avoid losing certain amount of tumor-related microglia subsets.

Considering the distinct roles of microglia and Mo/Mφ, it is indispensable to distinguish the two cell populations in myeloid cell researches in CNS pathologies. Nevertheless, known markers might be insufficient to stably differentiate these two populations during functional activation. As previously reported and confirmed in this study, during activation and differentiation, expression of signature genes for microglia (*P2ry12* and *Sall1*) and Mo/Mφ (*Ly6c* and *Ccr2*) were dramatically reduced (*19, 20, 46, 47*) (*7, 8*). On the other hand, Mo/Mφ were gradually assimilated after invading the CNS, considering the downregulation of CD45 and the upregulation of “microglia signature” genes such as *Cx3cr1* and *Tmem119*(*7, 8, 48*). Using the fate-mapping murine model and bulk-seq analysis, Bowman et al. reported that Itga4 can be used to distinguish microglia and Mo/Mφ(*7*). In the current study, *Crybb1* and *Ldhb* were identified as specific and stable markers across different microglia phenotypes in mice. Mo/Mφ clusters were featured with constant expression of *Iqgap1*, in keeping with multiple murine and clinical datasets. These newly-identified markers may assist the identification of the individual myeloid populations, and may facilitate a more precise immunotherapy of glioma and other CNS pathologies. On the basis of distinguishing the identity of microglia and Mo/Mφ, it is of greater biological significance to conduct comparative analysis between subsets from same ontology. For myeloid cells in clinical samples, directly comparing Mo/Mφ from tumor mass with microglia from normal tissues may confound the functional transition with identity difference. It is also worth noticing that different stages of tumor progression will inevitably lead to different compositions of myeloid subgroups. Therefore, we selected multiple time points to study the possible phenotypic and functional changes of myeloid cells.

Mo/Mφ were to blame for the reported pro-tumor phenotypes of myeloid cells. In addition to their high expression of genes related to angiogenesis and tumor invasion (ARG1, MMP14, VEGF-A), Mo/Mφ subsets were highly enriched in pathways related to regulation of T cell apoptotic process. This is consistent with our previous research that PD-L1 mainly distributed inside the tumor rather than periphery, suggesting that Mo/Mφ may be the main induction of PD-1^+^ T cell apoptosis (*49*). Meanwhile, abundance of Mo/Mφ subsets were negatively correlated with patient prognosis(*10, 12, 13*), and expression of Mo/Mφ related genes, ITGA4 and IQGAP1, were reported to show correlation with unfavorable prognosis (*7, 50*). Our study confirmed the prognostic significance of Mo/Mφ and found that this correlation was more significant in patients with high infiltration of peripheral immune cells. Recent studies demonstrated that targeting CCR2 to restrict Mo/Mφ extended the survival of glioma model mice, confirming the tumor-promoting role of Mo/Mφ (*51*). These results demonstrated that infiltration of Mo/Mφ might serve as a prognostic indicator for glioma patient and a promising target for glioma immunotherapy.

Glioma-associated microglia were featured with enhanced lipid metabolism, phagocytosis, and immune regulation at the terminal stage of transition. Notably, previous studies have observed similar characteristics of microglia in AD model mice and patients as well as cuprizone-fed mice and multiple sclerosis patients(*39, 52-54*). Collectively, these results indicated that microglia might evolve to present common disease-associated pattern, aiming to protect the neural tissue and restore the CNS homeostasis. In addition, “quiescent microglia score” was found positively corelated with survival time. Therefore, boosting the inflammatory response of microglia or eliminating the disease-associated microglia might either crush the CNS under overactivated neuroinflammatory response or weaken the CNS confronting physiological and pathological challenges. These findings may explain, at least in part, that no obvious clinical benefit was observed in the current immunotherapy targeting both microglia and Mo/Mφ(*2*).

Further studies are needed to clarify the role of microglia in glioma development. However, it is difficult to get direct evidence for this question. Existing methods, such as CSF1R inhibitors and DTR mice, may be insufficient to specifically and persistently deplete microglia, since the regeneration of residual microglia usually occurs within 1 month(*55*). Administration of CSF1R inhibitors after tumor establishment will inevitably disturb the development of tumor(*1, 56*). In addition, microglia and macrophages can also affect tumor via antigen-presentation to T cells. More specific research of myeloid cells-T cells-tumors axis is needed to completely reveal the functional characteristics of myeloid cells. Besides, other myeloid cells, such as dendritic cells, granulocytes, and mast cells, are not discussed in this study, and these cells may also play an important role in the process of glioma.

## Materials and Methods

### Experimental Design

This study aimed at characterizing the dynamics of CNS-resident microglia and the peripheral-derived Mo/Mφ, as well as comparing their functional differences during the development of glioma. For this, we used a combination of scRNA sequencing and microglial fate-mapping system. Most single-cell sequencing analysis mainly captured GAMs inside the tumor foci, while microglia actually gathered around the tumor. Therefore, existing datasets may not be sufficient to completely depict the dynamic changes of microglia. In this study, we sorted microglia and Mo/Mφ from tumor-bearing hemispheres at different stage of glioma, so as to reveal the authentic temporal and functional changes of these subsets. Flow cytometric analysis, in situ microscopy and prognostic analysis were then used to verify the computational analysis.

Experiments were performed with both males and females. For all mouse experiments, mice were randomly distributed into different groups. Numbers of animals and statistical analysis methods are given in each figure. Experiments (FACS, Immunofluorescence staining, and RNAscope assay) were repeated at least for three times and the representative results are shown. Catalog numbers and the description of antibodies used throughout the study can be found as an additional technical sheet (Table S6).

### Animals

C57BL/6J mice (6–8 weeks) were purchased from Shanghai Slac Laboratory Animal Co., Ltd. *Cx3cr1*^CreER^ mice (B6.129P2(C)-*Cx3cr1*^*tm2*.*1(cre/ERT2)Jung*^ /J, Stock No: 020940) and *Rosa-26-tdTomato* reporter mice (B6.Cg-*Gt*(*ROSA*)*26Sor*^*tm14(CAG-tdTomato)Hze*^/J, Stock No: 007914) were purchased from the Jackson Laboratory. Microglia reporter mice (*Cx3cr1*^*CreER*^: *Rosa26*-*tdTomato* mice) at 4 weeks old (either gender) were injected intraperitoneally with tamoxifen daily at 100 mg/kg body weight for 5 days. Tamoxifen (Sigma-Aldrich) was dissolved in corn oil (Sigma-Aldrich) at a concentration of 20 mg/ml. All mice were housed under the pathogen-free condition in the Animal Facility of Fudan University (Shanghai, China). All animal experiments adhered to the Guidelines for the Care and Use of Laboratory Animals and were approved by the Institutional Animal Care and Use Committee of Fudan University.

### Orthotopic glioma model

Microglia reporter mice or WT C57BL/6J mice were injected with GL261 glioma cells at 8 weeks old. Detailed procedure for stereotactic intracranial tumor inoculation was previously reported(*18*). Briefly, anesthetized mice were immobilized and mounted onto a stereotactic head holder in the flat-skull position. The skin of the skull was dissected in the midline by a scalpel. 1 mm anterior and 2 mm lateral to the bregma, the skull was carefully drilled with a 20-gauge needle tip. Then a microliter Hamilton syringe was inserted to a depth of 3 mm and retracted to a depth of 2.5 mm from the dural surface. 5 μL (2×10^4^ cells/μL) cell suspension or PBS was slowly injected in 2 min. The needle was then slowly taken out from the injection canal, and the skin was sutured.

### Immunofluorescence and RNAscope

Brains of orthotopic glioma model were harvested at day 10, 20, 30 after tumor inoculation. Frozen sections of mice brain were obtained as previously described(*18*). For immunofluorescence, sections were thawed and dried at room temperature and rinsed in PBS and followed by 4% PFA cell fixation (5 minutes at room temperature). Samples were permeabilized with 0.25% Triton-X 100 (15 minutes at room temperature) and blocked with blocking buffer containing 10% donkey serum (2 hours at room temperature or overnight at 4°C). Then, samples were incubated with indicated primary antibodies overnight at 4°C. Samples were then washed with PBS and incubated with the appropriate fluorophore-conjugated secondary antibodies: Alexa Flour-488, 594 (Thermo Fisher) at a dilution of 1:500 in 1% BSA for 1 hour at room temperature, and 4’, 6-diamidino-2phenylindole (DAPI) were used as a counterstain. For RNAscope, RNAscope Multiplex Fluorescent v2 Assay combined with Immunofluorescence was performed according to the manufacturer’s protocol (ACD Biosystems). IBA1 antibody followed with Alexa Flour-488-conjugated secondary antibody was used as the myeloid cell marker for each probe. Additionally, the ACD 3-plex negative control probe was used to confirm signal specificity. The probes (Probe-Mm-*Ccl3*-C2, Probe-Mm-*Mmp14*-C3, Probe-Mm-*Cst7*, and Probe-Mm-*Mki67*-C3) were amplified and labeled with Opal 570 or Opal 620 (Akoya Biosciences) for each experiment. Images were acquired by a fluorescence microscope Olympus IX73 (Olympus). Appropriate gain and black level settings were determined by control tissues stained with secondary antibodies alone. Analyses of images were performed using ImageJ software (U.S. National Institutes of Health).

### Fluorescence-activated cell sorting analyses

Percoll gradient isolation of immune cells were conducted as previously reported(*18*). Briefly, tumor-bearing hemispheres were harvested with olfactory bulbs and cerebella removed, dissociated mechanically with Dounce homogenizers to make homogenates and then forced through a filter for single-cell suspension and washed with PBS. Cell pallets were re-suspended in 37% Percoll. Percoll gradients (70%/37%/30%/0%) were set up and centrifuged for 20 min at 200 *g*, 4 °C, brake off. Mononuclear cells were collected at 70%/37% interface and washed with PBS. The majority of microglia located at the periphery of tumor foci. To avoid losing these microglial subsets, we thus isolated the myeloid cells from the tumor-bearing hemispheres, but not only the tumor mass. For flow cytometry, isolated cells were counted and incubated with anti-CD16/32 (eBioscience) for 30 minutes, followed by another 30-minute incubation with conjugated antibodies for extracellular markers. For intracellular cytokine detection, cells were stimulated in vitro with Cell Stimulation Cocktail (eBioscience) for 4 hours at 37°C before FACS analysis. After stimulation, the cells were stained for surface markers and cytokines with Intracellular Fixation and Permeabilization Buffer Set (eBioscience). Proper isotype controls and compensation controls were performed in parallel. BD FACSCanto II Cell Analyzer was used for flow cytometry, and a BD FACS Melody Cell Sorter was used for cell sorting. FlowJo software (Tree Star) was used for FACS data analysis. CD45^lo/hi^CD11b^+^ myeloid cells were sorted for scRNA-seq profiling. Each sample was pooled from 5 mice (tumor-bearing hemispheres) at Day 10, Day 20, and Day 30 after tumor inoculation. The cell viability and density of samples were checked before following procedures.

### RNA isolation and library preparation

The scRNA-Seq libraries were generated using the 10X Genomics Chromium Controller Instrument and Chromium Single Cell 3’V3.1 Reagent Kits (10X Genomics). Briefly, the density of cells was adjusted to 1000 cells/μL. For each sample, approximately 7,000 cells were loaded into each channel to generate single-cell Gel Bead-In-Emulsions (GEMs), resulting into expected mRNA barcoding of 5,000 single-cells. After the RT step, GEMs were broken and barcoded-cDNA was purified and amplified. The amplified barcoded cDNA was fragmented, A-tailed, ligated with PCR amplified adaptors and index. The final libraries were quantified using the Qubit High Sensitivity DNA assay (Thermo Fisher Scientific) and the size distribution of the libraries were determined using a High Sensitivity DNA chip on a Bioanalyzer 2200 (Agilent). Then, all libraries were sequenced by illumina sequencer (Illumina) on a 150 bp paired-end run.

### Single-cell RNA data processing

We applied fastp with default parameter filtering the adaptor sequence and removed the low-quality reads. Using CellRanger v3.1.0, the feature-barcode matrices were obtained by aligning reads to the mouse genome (Mm10 Ensemble version 92). Finally, we applied the down sample analysis among sequenced samples according to the mapped barcoded reads per cell of each sample and achieved the aggregated matrix. Cell quality filtering removed cells containing over 200 expressed genes and with mitochondria UMI rate below 20%. Mitochondria genes were removed in the expression table. A total of 18453 single-cell RNA profiles was acquired.

### Unsupervised clustering and DEG identification

Seurat package (v2.3.4)(*57*) was used for cell normalization and scaled according to the UMI counts of each sample and percent of mitochondria rate. PCA was constructed based on the scaled data with top 2000 high variable genes and top 10 principal components were used for tSNE and UMAP construction. Utilizing graph-based cluster method, we acquired the unsupervised cell clustering result based on the top 10 principal components. Marker genes were calculated by FindAllMarkers function (test.use = “wilcox”, min.pct = 0.25, min.diff.pct = 0.1, logfc.threshold = 0.25). In order to identify the detailed cell clusters, clusters of same cell type were selected for further graph-based clustering and differential gene expression analysis. After first round clustering, cells not in the study scope were excluded, including NK cells, T cells, mast cells, and neutrophils. Finally, 15,765 single-cell RNA profiles were used in following analyses.

After cell identity annotation, we used the FindMarkers function (test.use = “wilcox”, min.pct = 0.25, min.diff.pct = 0.1) based on normalized data to identify DEGs. P -value adjustment was performed using Bonferroni correction based on the total number of genes in the dataset. DEGs with adjusted P -values >= 0.05 were filtered out. Genes of top 100 average Log2 fold change were recognized as the marker genes for microglia or Mo/Mφ population (Table S2). Genes of top 20% or 25% absolute average Log2 fold change were recognized as the up-regulated or down-regulated genes for each cluster. (Table S2).

### Definition of cell scores and identity

To clarify the identity of clusters, we concluded the microglia markers (P2ry12, Siglech, Sparc, Gpr34, Tmem119, Fcrls and Olfml3) and Mo/Mφ markers (Itga4, S100a6, S100a4, Ccr2, Tgfbi, Plac8 and Ly6c2)(*6-8, 19, 20*). Gene Set Variation Analysis (GSVA) (v1.38.0)(*21*) were applied to calculate “Microglia score” and “Mo/Mφ score”. Difference of z-score normalized microglia and Mo/Mφ score was calculated and recognized as myeloid identity score. Mixsmsn package (v1.1-8)(*58*) was used to fit finite mixture of scale mixture of skew-normal (FM-SMSN) distributions for myeloid identity score and 0.27 was selected as the cut-off for determining microglia and Mo/Mφ identity. Analysis of myeloid cells from other scRNA-seq datasets

We used the Seurat R package to perform normalization for the datasets of myeloid cells in various diseases (GSE98969, GSE121654, GSE103334, and GSE118948). We performed PCA on the normalized expression matrix using top 2000 highly variable genes identified by “FindVariableGenes” function. Following the results of PCA, the top 20 PCs were selected for clustering with the specific resolution parameters. We concluded the marker genes for disease-associated microglia (DAM) from the original published paper (Table S4). Gene Set Variation Analysis (GSVA) were applied to calculate “DAM score” and “MG4 score” based on the selected marker genes for DAM and MG4. Cluster with highest DAM score were recognized as DAM.

### Cell trajectory analysis

The cell trajectory of microglia and Mo/Mφ populations was inferred respectively by using Monocle2(*59*). We first created an object with parameter ‘‘expressionFamily = negbinomial.size’’ following the Monocle2 tutorial. Then we select marker genes of the Seurat clustering result to order the cells in trajectory analysis. Finally, trajectory analysis was performed using DDR-Tree and default parameter.

### Gene Enrichment Analysis and SCENIC Analysis

To characterize the relative activation of a given gene set, we performed GSVA analysis using standard settings. The gene sets were selected from the Gene Ontology (GO) (biological processes)(*60*) and Kyoto Encyclopedia of Genes and Genomes (KEGG) pathway database(*61*). To assess transcription factor regulation strength, we applied the Single-cell regulatory network inference and clustering (pySCENIC, v0.9.5)(*62*) workflow using the 20-thousand motifs database for RcisTarget and GRNboost.

### TCGA LGG/GBM data downloading and survival analyses

The normalized expression and phenotype data of TCGA LGG/GBM dataset(*63*) were downloaded from the GlioVis database(*64*). We excluded patients whose survival and status were not available. Cox proportional hazards model was used for survival analysis and Kaplan–Meier method for survival curves. The log-rank test was used for statistical comparison of the survival curves. R packages survival (v3.2-7) and survminer (v0.4.8) were applied for the analysis and plotting.

### Deconvolution of the cellular composition with MCP-counter

The MCP-counter package (v1.2.0)(*65*) was applied to produce the absolute abundance scores for eight major immune cell types (T cells, CD8^+^ T cells, cytotoxic lymphocytes, natural killer cells, B lineage cells, monocytic lineage cells, myeloid dendritic cells and neutrophils), endothelial cells and fibroblasts. The abundance scores of myeloid clusters were calculated based on reference genes (Table S4). The infiltration of peripheral immune cell was calculated by aggregating the z-score normalized abundance score of major immune cell types. Patients were evenly separated into high, median and low infiltration group based on the aggregated score. Dominant myeloid cells were determined based on the mean abundance score of Mono-Mo2-*Sell* and Mφ-MP2-*Mrc1* cluster minus abundance score of MG-MG0-*P2ry12* cluster. Mixtools package (v1.2.0)(*66*) was used to fit finite mixture of normal distributions for the calculated score and 0.35 was selected as the cut-off for microglia-dominant and monocyte/macrophage-dominant.

### Statistical Analysis

GraphPad Prism 9.0 (GraphPad Software Inc., USA) was used for FACS and IF data analysis. Parametric data were presented as mean ± standard error of the mean (SEM). Differences between two groups were analyzed using the Student’s unpaired t test. Analysis of variance (ANOVA) was used to compare multiple groups. Statistical significance was determined at *P* < 0.05.

## Acknowledgments

We thank Prof. Ji-Yang Wang and Dr. Ermeng Xiong for critiquing the article and helpful discussions. We thank Dr. Feizhen Wu for the support of bioinformatics analysis. We thank NovelBio Co., Ltd for with their NovelBrain Cloud Analysis Platform (www.novelbrain.com). We thank the support from the Innovative Research Team of High-level Local Universities in Shanghai.

## Author contributions

Conceptualization: JQ;

Methodology: JQ, BW, CL, CW;

Investigation: JQ, CW, CL;

Visualization: JQ, CW, CL;

Funding acquisition: YC, JQ;

Software: CW;

Supervision: YC;

Writing – original draft: JQ, CW;

Writing – review & editing: BW, KF, CZ, JL, ML, YL, YC

## Data and materials availability

The datasets generated during this study are available with accession number GSE171081 (The dataset will be publicly available after publication). The original data of myeloid cells in 5xFAD mice, Aging and LPC-injected mice, CK-p25 mice and EAE mice for Fig. 5c in this paper are available with accession number GSE98969, GSE121654, GSE103334, and GSE118948 respectively(*39-42*). All of the code will be available on a github repository after publication. All other data are available upon reasonable request.

## Supplementary Materials

Fig. S1. Validation of reporter mice and gating strategy for myeloid cells.

Fig. S2. Expression of cluster markers.

Fig. S3. Expression of marker genes for pan-myeloid cells, microglia and Mo/Mφ.

Fig. S4. Single-cell profiling of myeloid cells in human glioma sample.

Fig. S5. Signaling and metabolic pathway enrichment of myeloid cells in human glioma sample.

Fig. S6. SCENIC analysis of myeloid cells.

Fig. S7. Survival analysis based on the expression of microglia and Mo/Mφ markers.

Table S1. Absolute cell counts of each cell population at different time points. Figure 2 related.

Table S2. Featured genes of microglia and Mo/Mφ clusters. Figure 3 and Figure 4 related.

Table S3. GO and KEGG pathways. Figure 5 and Figure 6 related.

Table S4. Marker genes for disease-associated microglia. Figure 6 related.

Table S5. Marker genes of each cell population used in cellular composition analysis with MCP-counter. Figure 7 related.

Table S6. Antibodies used in this study.

