## Supplementary Material for "Dynamics of glioma-associated microglia and macrophages reveals their divergent roles in the immune response of brain"

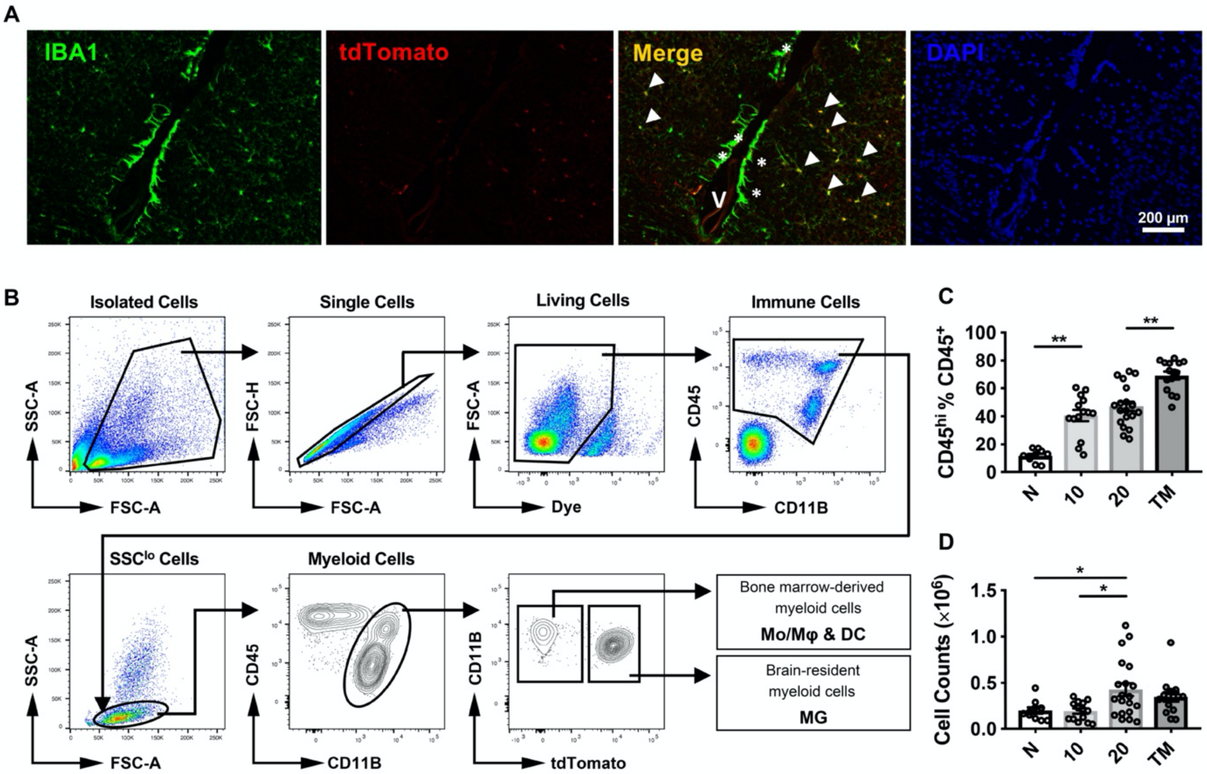


**Fig. S1. Validation of reporter mice and gating strategy for myeloid cells.**

(**A**) Representative IF staining of normal brain tissue from microglial reporter mice. IBA1 (green), tdTomato (red) and DAPI (blue). V, lateral ventricle. Arrows, $IBA1^{+}$ $tdTomato^{+}$ microglia. Asterisk, $IBA1^{+}$ $tdTomato^{-}$ intraventricular macrophages (peripheral derived myeloid cells). Scale bar, 200 µm.

(**B**) Gating strategy of flow cytometric analysis for myeloid cells, microglia and Mo/Mφ.

(**C**) The statistical summary for abundance of the myeloid populations, n = 10-20. Data pooled from 4 independent experiments. (**D**) the absolute cell counts of the myeloid populations isolated from normal or tumor-bearing hemispheres, n = 10-20. Data pooled from 4 independent experiments. One-way ANOVA was performed in C and D. $^{*}P < 0.05$; $^{**}P < 0.01$. All values are shown as mean ± SEM.

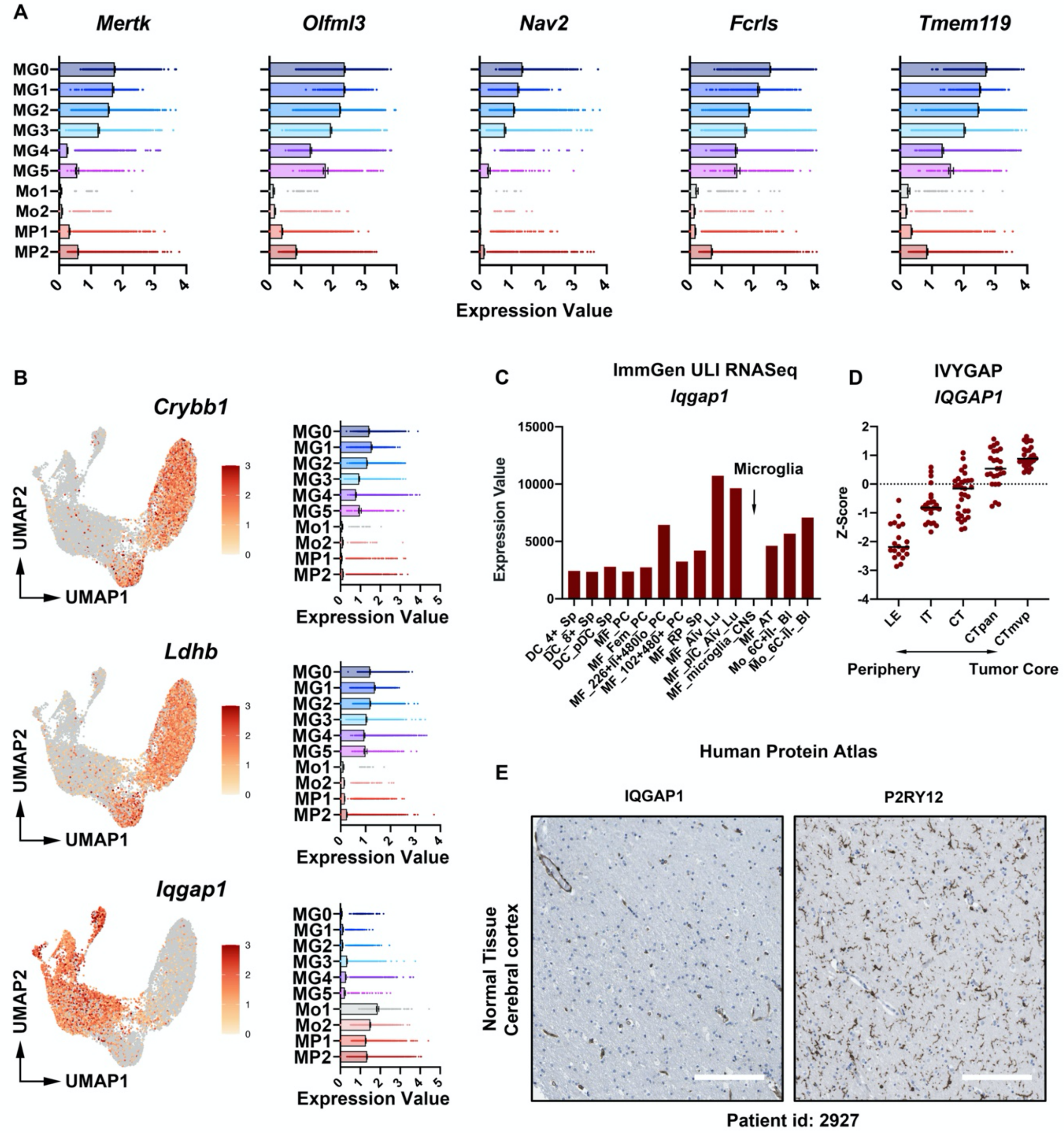


**Fig. S2. Expression of cluster markers.**

(**A**) Bar plots showing the upregulation of microglia marker genes in Mo/Mφ.

(**B**) Left, UMAP projection showing the expression of Crybb1, Ldhb and Iqgap1. Right, Bar plots showing the expression of genes in each microglia and Mo/Mφ clusters.

(**C**) Iqgap1 expression of different murine myeloid cell subsets. Data from Immunological Genome Project (ImmGen).

(**D**) IQGAP1 expression at different areas of human glioma samples, n = 20-31. LE, leading edge; IT, infiltrating tumor; CT, cellular tumor; CTpan, cellular tumor-pseudopalisading cells around necrosis; CTmvp, cellular tumor-microvascular proliferation. Data from Ivy Glioblastoma Atlas Project (IVYGAP).

(**E**) Immunohistochemistry staining for IQGAP1 and P2RY12 in human normal brain. Image credit, Human Protein Atlas. Scale bar, 200 mm.

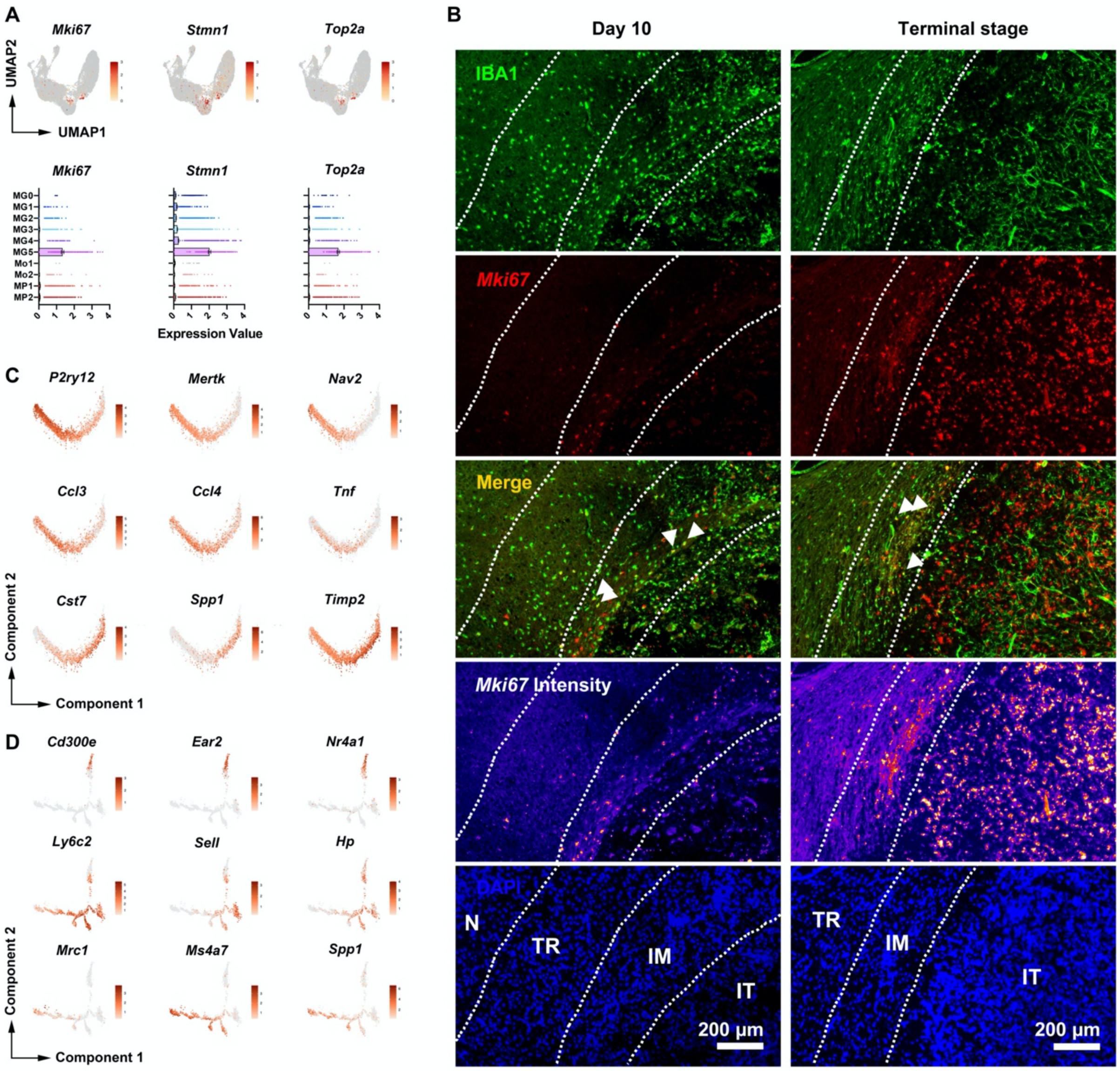


**Fig. S3. Expression of marker genes for pan-myeloid cells, microglia and Mo/Mφ.**
(**A**) UMAP projection (top) and bar plots (bottom) showing the expression of proliferation-associated genes.
(**B**) Representative figure of RNAscope staining in murine glioma. Representative staining for IBA1 (green), Mki67 (red) and DAPI (blue) in the tumor-bearing hemisphere at Day 10 and Terminal Stage. N, normal tissue; TR, tumor rim; IM, invasive margin; IT, intratumoral region. Dashed line, border of different areas. Arrow, Mki67+ myeloid cells. Scale bar, 200 μm.
(**C**) 2D pseudotime plots showing the dynamics of marker genes in microglia-MG0-P2ry12 (top), microglia-MG1-Ccl3/microglia-MG2-Ccl3 (middle) and microglia-MG3- Csf1/microglia-MG4-Cst7 (bottom).
(**D**) 2D pseudotime plots showing the dynamics of marker genes in monocyte-Mo1-Nr4a1 (top), monocyte-Mo2-Sell/macrophage-MP1Vegfa (middle) and macrophage-MP2-Mrc1 (bottom).

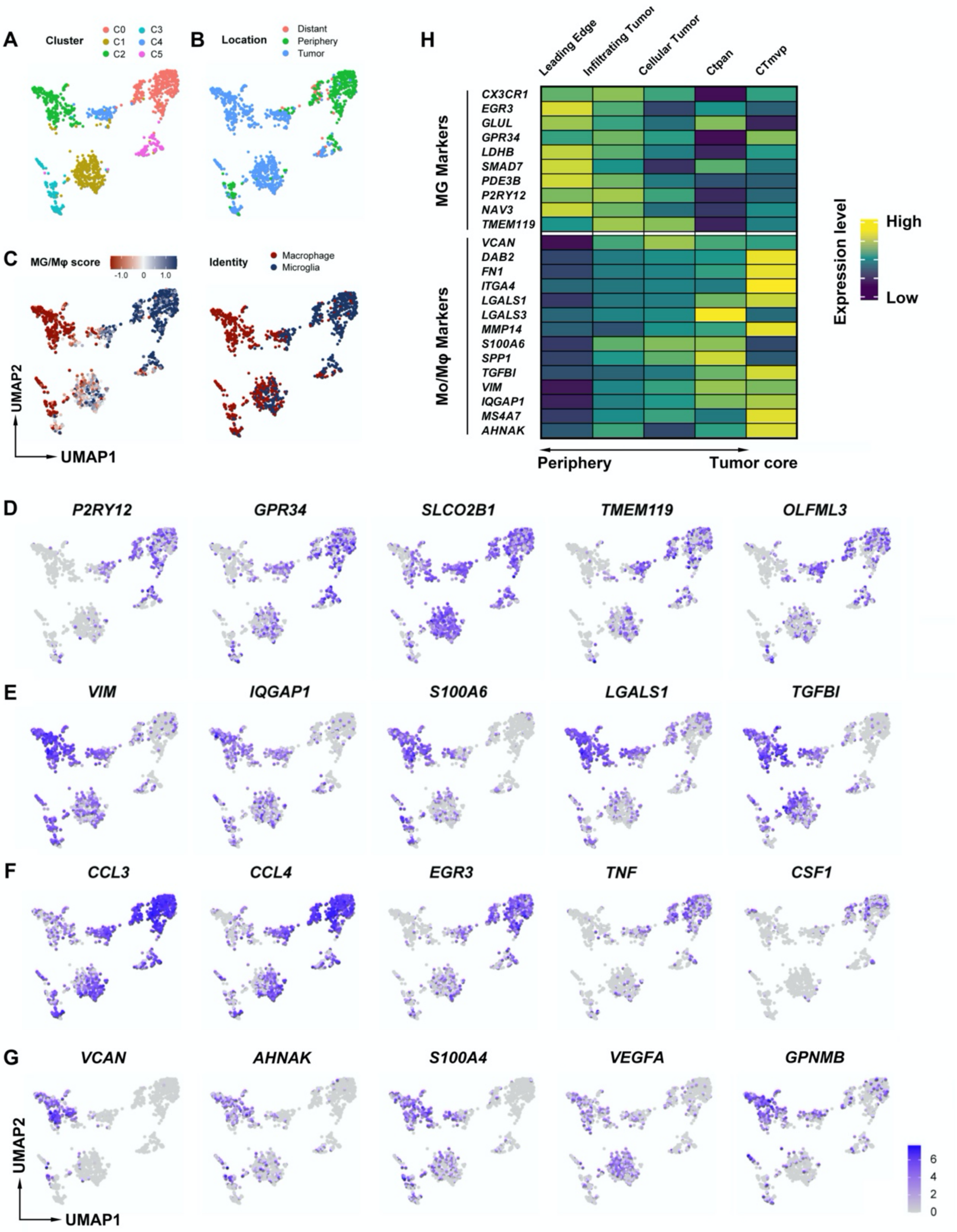


**Fig. S4. Single-cell profiling of myeloid cells in human glioma sample.**

(**A-B**) UMAP projection showing the single-cell profiling (**A**) and location (**B**) of 1,415 myeloid cells from human glioma sample.

(**C**) UMAP projection showing the identity score (left) and identity of myeloid cells (right). GSVA was applied to calculate the identity score based on the expression of marker genes.

(**D-G**) UMAP projection showing the expression of microglia markers (**D**), Mo/Mφ markers (**E**), function-related genes in microglia (**F**), and function-related genes in Mo/Mφ (**G**).
(**H**) Heatmap showing the expression of microglia and Mo/Mφ markers at different areas of human glioma samples. LE, leading edge; IT, infiltrating tumor; CT, cellular tumor; CTpan, cellular tumorpseudopalisading cells around necrosis; CTmvp, cellular tumor-microvascular proliferation. Data from Ivy Glioblastoma Atlas Project. Data from GSE84465.

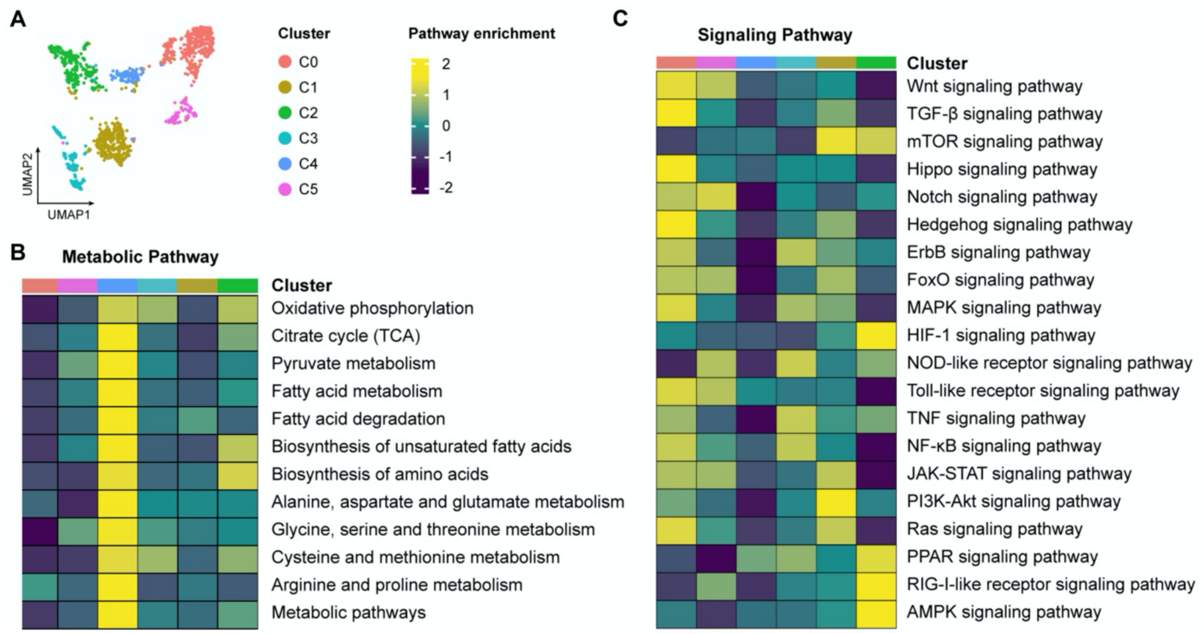


**Fig. S5. Signaling and metabolic pathway enrichment of myeloid cells in human glioma sample.**

(**A**) UMAP projection showing the single-cell profiling of 1,415 myeloid cells from human glioma sample (data from GSE84465).

(**B-C**) Heatmap indicating the enrichment of metabolic (**B**) and signaling (**C**) pathways in myeloid cells from human glioma sample (data from GSE84465). GSVA was applied to calculate the pathway enrichment based on the mean expression of genes within each subset.

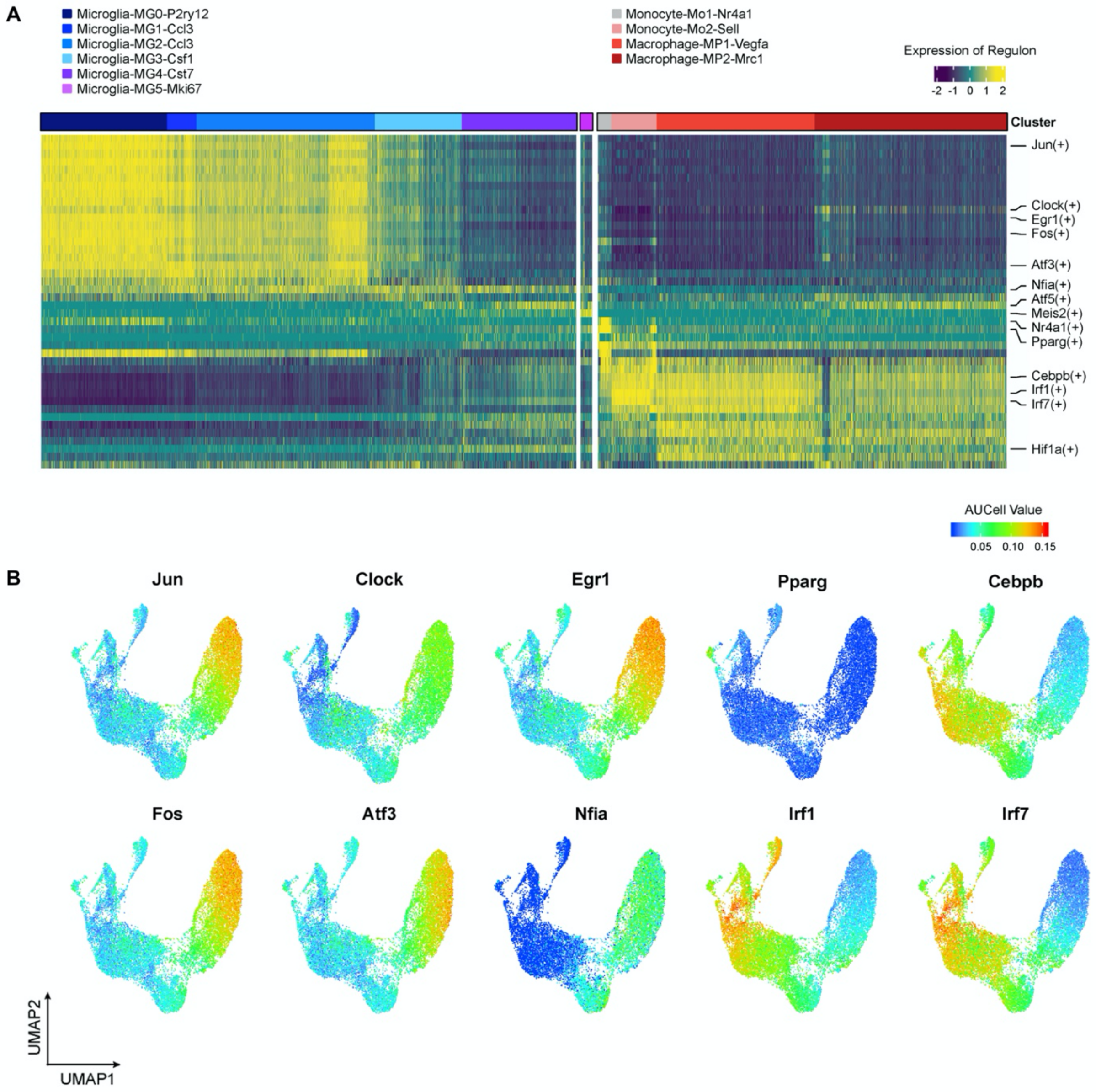


**Fig. S6. SCENIC analysis of myeloid cells.**

(**A**) Heatmap indicating the regulon activity of microglia and Mo/Mφ populations. (**B**) UMAP projection showing the AUCell value of featured regulons in myeloid cell populations.

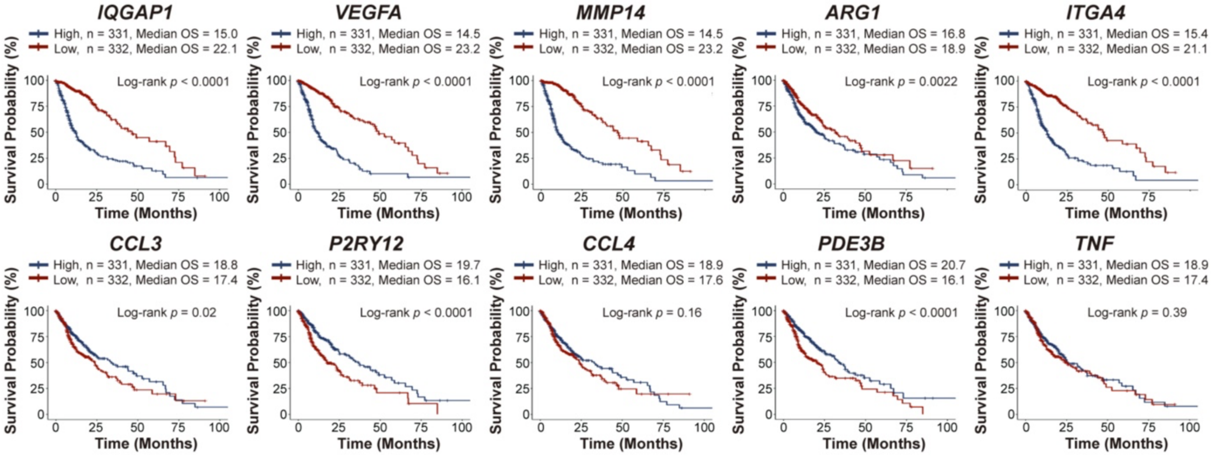


**Fig. S7. Survival analysis based on the expression of microglia and Mo/Mφ markers.** Kaplan-Meier analysis showing the survival probability of patients, characterized by either high (blue) or low (red) expression of Mo/Mφ (top) and microglia (bottom) marker genes.

**Table S1. Absolute cell counts of each cell population at different time points.** Fig. 2 related.

| | Day 10 | %(Day 10) | Day 20 | %(Day 20) | Terminal Stage | %(Terminal Stage) |
|---|---|---|---|---|---|---|
| **DC-DC1-*Ciita*** | 227 | 5.58 | 236 | 5.33 | 613 | 8.43 |
| **DC-DC2-*Ccr7*** | 143 | 3.52 | 169 | 3.82 | 313 | 4.30 |
| **Microglia-MG0-*P2ry12*** | 579 | 14.24 | 1141 | 25.79 | 136 | 1.87 |
| **Microglia-MG1-*Ccl3*** | 263 | 6.47 | 159 | 3.59 | 12 | 0.16 |
| **Microglia-MG2-*Ccl3*** | 1330 | 32.71 | 855 | 19.32 | 426 | 5.86 |
| **Microglia-MG3-*Csf1*** | 381 | 9.37 | 320 | 7.23 | 575 | 7.90 |
| **Microglia-MG4-*Cst7*** | 354 | 8.71 | 398 | 8.99 | 936 | 12.87 |
| **Microglia-MG5-*Mki67*** | 55 | 1.35 | 46 | 1.04 | 86 | 1.18 |
| **Monocyte-Mo1-*Nr4a1*** | 83 | 2.04 | 74 | 1.67 | 53 | 0.73 |
| **Monocyte-Mo2-*Sell*** | 156 | 3.84 | 194 | 4.38 | 313 | 4.30 |
| **Macrophage-MP1-*Vegfa*** | 334 | 8.21 | 514 | 11.62 | 1469 | 20.20 |
| **Macrophage-MP2-*Mrc1*** | 161 | 3.96 | 319 | 7.21 | 2342 | 32.20 |
| **Total** | 4066 | | 4425 | | 7274 | |

**Table S2. Featured genes of microglia and Mo/Mφ clusters.** Fig. 3 and Fig. 4 related. (Submitted as excel file named: Table S2. xlsx) (separate file)

**Table S3. GO and KEGG pathways.** Fig. 5 and Fig. 6 related. (Submitted as excel file named: Table S3. xlsx) (separate file)

**Table S4. Marker genes for disease-associated microglia.** Fig. 6 related.

| Model | GEO | Gene for DAM | Gene for MG4 (GSE171081) |
|---|---|---|---|
| 5xFAD | GSE98969 | Cst7, Itgax, Csf1, Lpl, Clec7a, Axl, Trem2, Cd9, Ctsl, Ccl6, Lilrb4, Timp2 | Cst7, Cd72, Fabp5, Spp1, Mif, Cstb, AW112010, Ccl5, Irf7, Lgals3, Fxyd5 |
| Aging | GSE121654 | Ccl4, Il1b, Lpl, Fam20c, Cst7, Csf1, Ifitm3, Ifit3, Rtp4, Irf7, Isg15, Oasl2 | |
| CK-p25 | GSE103334 | C3, C4b, Cfb, Axl, Clec7a, Lgals3, H2-D1, H2-Q5, H2-Aa, H2-Ab1, Cd74, Ifitm3, Irf7, Oas1a, Rsad2, Zbp1 | |
| EAE | GSE118948 | Ly86, Ccl5, Apoe, Cd74, Cxcl10, Ccl2, Mki67 | |
| LPC | GSE121654 | Ccl12, Cst7, Ifi204, Axl, Lpl | |

**Table S5. Marker genes of each cell population used in cellular composition analysis with MCP-counter.** Fig. 7 related.

| Cell population | Gene |
|---|---|
| Microglia | P2RY12, LDHB, SPARC, GPR34, HPGDS, LAG3, CFH, MERTK, SLCO2B1, TMEM119, GLUL, LTC4S, PLXDC2, OLFML3, PTGS1, PDE3B, CRYBB1, CD81 |
| BMDM | VIM, IQGAP1, PLBD1, ITGA4, IL1B, S100A6, LGALS1, IFITM2, S100A4, CYBB, S100A11, TGFBI, CRIP1, LSP1 |
| Mono-Mo2-*SELL* | S100A6, S100A4, ANXA2, AHNAK, S100A10, IFITM1, ISG20, SELL, MXD1 |
| MG-MG0-P2RY12 | TMEM119, P2RY12, NAV3, SAMD7, GLUL, PDE3B, GPR34 |
| Mφ-MP2-MRC1 | TMEM119, SPP1, DAB2, MS4A7, MMP14, ARG1 |
| T cells | CD28, CD3D, CD3G, CD5, CD6, CHRM3-AS2, CTLA4, FLT3LG, ICOS, MAL, MGC40069, PBX4, SIRPG, THEMIS, TNFRSF25, TRAT1 |
| CD8 T cells | CD8B |
| Cytotoxic lymphocytes | CD8A |
| B lineage | BANK1, CD19, CD22, CD79A, CR2, FCRL2, IGKC, MS4A1, PAX5 |
| NK cells | CD160, KIR2DL1, KIR2DL3, KIR2DL4, KIR3DL1, KIR3DS1, NCR1, PTGDR, SH2D1B |
| Monocytic lineage | ADAP2, CSF1R, FPR3, KYNU, PLA2G7, RASSF4, TFEC |
| Myeloid dendritic cells | CD1A, CD1B, CD1E, CLEC10A, CLIC2, WFDC21P |
| Neutrophils | CA4, CEACAM3, CXCR1, CXCR2, CYP4F3, FCGR3B, HAL, KCNJ15, MEGF9, SLC25A37, STEAP4, TECPR2, TLE3, TNFRSF10C, VNN3 |
| Endothelial cells | ACVRL1, APLN, BCL6B, BMP6, BMX, CDH5, CLEC14A, DIPK2B, EDN1, ADGRL4, EMCN, ESAM, ESM1, FAM124B, HECW2, HHIP, KDR, MMRN1, MMRN2, MYCT1, PALMD, PEAR1, PGF, PLXNA2, PTPRB, ROBO4, C, SHANK3, SHE, TEK, TIE1, VEPH1, VWF |
| Fibroblasts | COL1A1, COL3A1, COL6A1, COL6A2, DCN, GREM1, PAMR1, TAGLN |

**Table S6. Antibodies used in this study.**

| **FACS Antibodies** | | | | |
|---|---|---|---|---|
| **Antigen** | **Conjugation** | **Clone** | **Source** | **Cat#** |
| CD45 | eFluor 450 | 30-F11 | Thermofisher | 48-0451-82 |
| CD45 | PE-Cyanine7 | 30-F11 | Thermofisher | 25-0451-82 |
| CD11b | FITC | M1/70 | Thermofisher | 11-0112-41 |
| CD11b | PE-Cyanine7 | M1/70 | Thermofisher | 25-0112-82 |
| Ly6C | PE-Cyanine7 | HK1.4 | Thermofisher | 25-5932-82 |
| P2RY12 | APC | S16007D | Biolegend | 848006 |
| CCL3 | eFluor 660 | DNT3CC | Thermofisher | 50-7532-82 |
| TNF-a | APC | MP6-XT22 | Thermofisher | 17-7321-82 |
| ARG1 | PE-Cyanine7 | A1exF5 | Thermofisher | 25-3697-82 |
| VEGF-A | / | SP07-01 | Thermofisher | MA5-32038 |
| PD-L1 | PE-Cyanine7 | MIH5 | Thermofisher | 25-5982-82 |
| MHC-II | PerCP-eFluor710 | AF6-120.1 | Thermofisher | 46-5320-82 |
| **IF Antibodies** | | | | |
| **Antigen** | **Host** | **Clone** | **Source** | **Cat#** |
| Antigen/Host/Clone/Supplier/Cat.No. | | | | |
| IBA1 | rabbit | Polyclonal | Wako | 019-19741 |
| P2RY12 | rabbit | Polyclonal | Anaspec | AS-55043A |
| ARG1 | rat | A1exF5 | Thermofisher | 25-3697-82 |
| **IF Second Antibodies** | | | | |
| **Antigen** | **Host** | **Conjugation** | **Source** | **Cat#** |
| rabbit IgG | donkey | Alexa Fluor 488 | Thermofisher | A-21206 |
| rat IgG | donkey | Alexa Fluor 594 | Thermofisher | A-21206 |
